# Universal Loop assembly (uLoop): open, efficient, and species-agnostic DNA fabrication

**DOI:** 10.1101/744854

**Authors:** Bernardo Pollak, Tamara Matute, Isaac Nuñez, Ariel Cerda, Constanza Lopez, Valentina Vargas, Anton Kan, Vincent Bielinski, Peter von Dassow, Chris L. Dupont, Fernán Federici

## Abstract

Standardised Type IIS DNA assembly methods are becoming essential for biological engineering and research. Although a ‘common syntax’ has been proposed to enable higher interoperability between DNA libraries, Golden Gate (GG)-based assembly systems remain specific to target organisms. Furthermore, these GG assembly systems become laborious and unnecessarily complicated beyond the assembly of 4 transcriptional units. Here, we describe “universal Loop” (uLoop) assembly, a simple system based on Loop assembly that enables hierarchical fabrication of large DNA constructs (> 30 kb) for any organism of choice. uLoop comprises two sets of four plasmids that are iteratively used as odd and even levels to compile DNA elements in an exponential manner (4^n-1^). The elements required for transformation/maintenance in target organisms are also assembled as standardised parts, enabling customisation of host-specific plasmids. Thus, this species-agnostic method decouples efficiency of assembly from the stability of vectors in the target organism. As a proof-of-concept, we show the engineering of multi-gene expression vectors in diatoms, yeast, plants and bacteria. These resources will become available through the OpenMTA for unrestricted sharing and open-access.

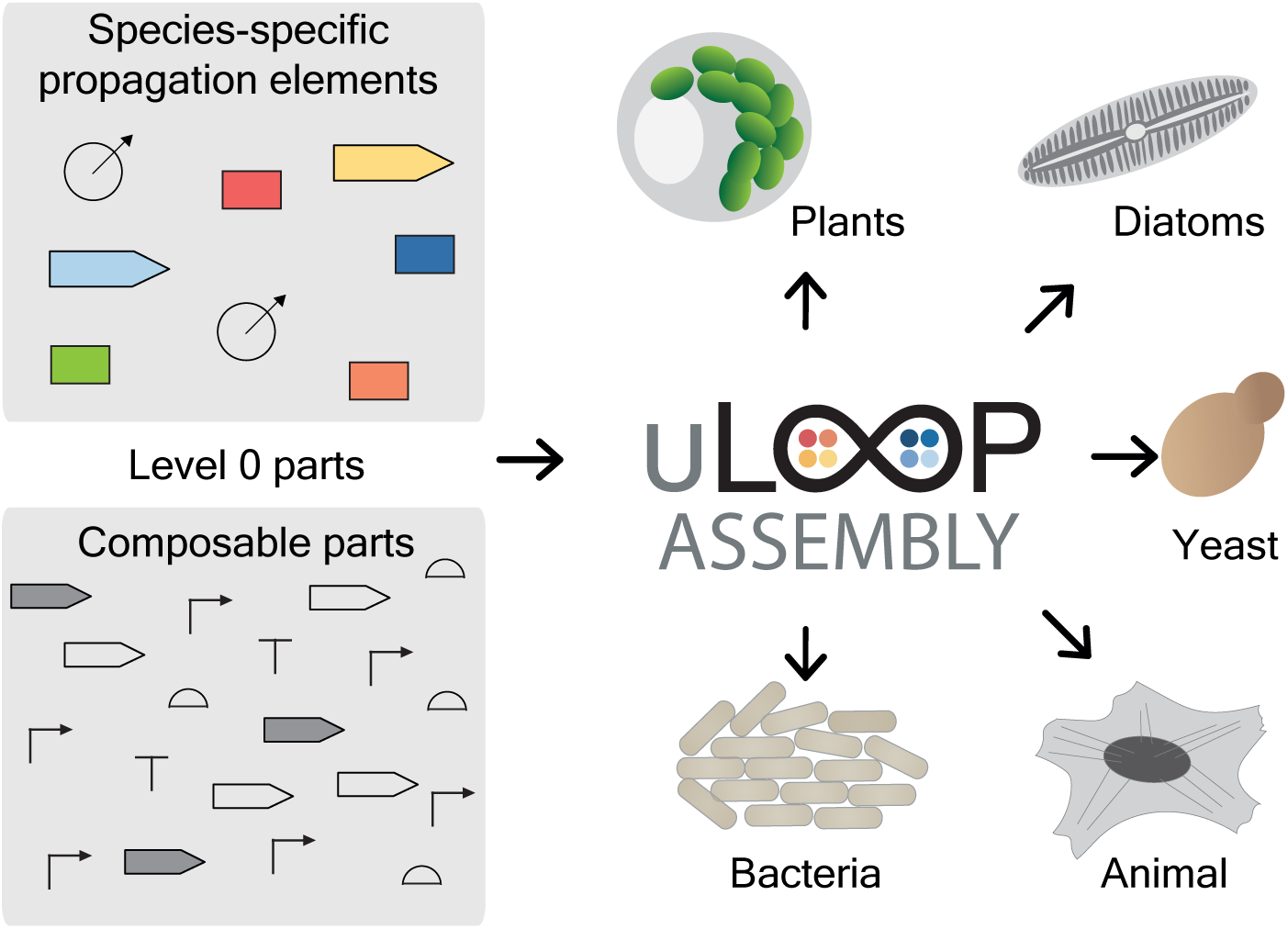

## INTRODUCTION

Methods for DNA assembly have been instrumental in the pursuit of understanding the genetic code and our capacity to engineer biological functions. Standardised methods for DNA assembly have gained popularity in recent years, due to their accessibility, reliability and amenability to automation (1–3). Among these, Type IIS restriction endonuclease-based assembly systems have been adopted in different fields of biology due to their high efficiency and versatility (4–12).

The development of Type IIS assembly systems has enabled the establishment of a ‘common syntax’ that allows basal DNA parts from different Type IIS assembly systems to be interoperable (8,13). This has led to a vast library of domesticated DNA parts available for use. As a consequence of the increased need for sharing and exchange, the universal biological material transfer agreement has been revised (14) and an open material transfer agreement is being proposed to facilitate and expedite the sharing and exchange of plasmids (15). Further, the need for IP-free DNA assembly systems to enable open access to DNA fabrication has been addressed, as in the case of the Loop assembly method developed previously by the authors (10).

Loop assembly is a recursive Type IIS DNA assembly system that enables reliable assembly of DNA fragments of exponentially increasing size. The use of inverted orientations of BsaI and SapI recognition sites in receiver vectors for odd and even levels of assembly (i.e. L1, L2, L3…) allows the product of one level to become the substrate for following level. This design permits the use of a compact number of plasmids (two sets of four odd and even vectors), which are utilised repeatedly in alternating steps. Thus, Loop assembly provides a simple system for the parallel composition of 4 genetic modules per vector and assembly level. Despite its efficiency and simplicity, Loop assembly was originally developed in binary vector backbones used for *Agrobacterium*-mediated transformation of plants, thus its use has been largely restricted to plant biology. Similarly, most Type IIS assembly systems have been restricted to particular organisms or taxonomic groups due to the presence of host-specific elements in the plasmid backbones required for their propagation in the host; e.g. yeast (16), plants (6,11), mammalian (17) and microalgae (18).

Historically, elements required for transformation, conjugation and transfection of DNA into organisms of choice have been treated as integral components of plasmid backbones used for cloning and assembly. These elements are refactored from natural systems, such as the T-DNA from the tumor-inducing (Ti) plasmid of *Agrobacterium tumefasciens*, the origin of transfer (oriT) from the RK2 plasmid for bacterial conjugation, or autonomous replicating sequences (ARS) that enable extrachromosomal replication of plasmids in some eukaryotes. While well-established model systems such as yeast and plants can be manipulated with these engineered vectors from seminal work, new model organisms often require the development of new vector systems. For example, diatom transformation was implemented by the combination of conjugation machinery from the RK2 plasmid and ARS from yeast in order to enable bacterial conjugation and posterior replication of the conjugated DNA as an extrachromosomal episome (19). The use and reuse of plasmids is not generally possible without significant alteration due to the lack of modularity and standards in plasmid engineering.

Thus, elements required for transformation into organisms of choice and sequences designed for cloning and plasmid assembly have remained coupled in DNA assembly systems. While no distinction has been made between these two plasmid functions, now it is possible to address them separately. Decoupling of the DNA assembly logic from elements required for DNA transfer and maintenance in the organism of choice can yield more compact vectors, which can be engineered and characterised for efficient multipart assembly in *Escherichia coli*. Elements required for transformation can be modularised and included during the assembly routine alongside with other transcription units (TUs) to generate species-specific plasmids capable of transfer and propagation in the organism of interest. This strategy would enable the use of the same assembly vectors in different organisms, already engineered and characterised for multi-part assembly.

Here, we describe Universal Loop assembly (uLoop), a system for open, efficient and species-agnostic DNA fabrication. We have decoupled the assembly logic of Loop assembly from the host-specific propagation elements to enable universal DNA assembly that retains high efficiency regardless of the final host. We utilised 3 vectors broadly used in the synthetic biology community as well as the pCAMBIA vector (https://cambia.org/) in order to implement the Loop assembly logic (10). In this strategy, elements for propagation in the host are included in the assembly routine for their subsequent use in the organism of interest. We demonstrate the capability of uLoop to generate vectors containing elements for conjugation and show their function through multi-spectral fluorescent protein expression in the diatom *Phaeodactylum tricornutum*, the yeast *Saccharomyces cerevisiae*, protoplasts of *Arabidopsis thaliana* and in *Escherichia coli*.

## MATERIAL AND METHODS

### Plasmid construction

The uLoop vector kits were generated using Gibson assembly (20). First, vectors pCAMBIA, pJT170 (21), pSB4K5 (22) and pAN3945 (23) were re-annotated to determine the position of the ORI and the antibiotic resistance cassette. Then, additional elements not related to basic plasmid propagation were removed *in silico*, with the exception of pSB4K5 where prefix and suffix flanking terminator sequences were kept, and pJT170 where the *rop* gene was maintained. Sequences were screened for BsaI and SapI sites and then assemblies were designed to remove these restriction sites and other non-essential elements. Resulting plasmids were then used to generate a secondary version with an alternate antibiotic resistance marker (spectinomycin resistance for pCA and pCO vectors in even levels; chloramphenicol for pSB and pAN vectors in even levels). Finally, the Loop assembly schema was introduced into each vector kit using gBlocks through Gibson assembly, generating the uLoop vector kits. DNA parts were domesticated into L0 using the pL0R-lacZ vector, an entry vector generated to domesticate sequences using SapI.

### Type IIS assembly

The Loop Type IIS assembly protocol was adapted from Pollak et al., 2019 (10), and can be found at https://www.protocols.io/view/loop-and-uloop-assembly-yxnfxme. An aliquot of 15 fmol of each part to be assembled was mixed with 7.5 fmol of the receiver plasmid in a final volume of 5 µl with distilled H_2_O (dH_2_O). The reaction master mix for odd level reactions was prepared using 3 µl of dH_2_O, 1 µl of T4 DNA ligase buffer 10X (NEB cat. B0202), 0.5 µl of 1 mg mL^−1^ purified bovine serum albumin (1:20 dilution in dH_2_O of BSA, Molecular Biology Grade 20 mg ml^−1^, NEB cat. B9000), 0.25 µl of T4 DNA ligase at 400 U µL^−1^ (NEB cat. M0202) and 0.25 µl of BsaI (NEB cat. R0535) at 10 U µL^−1^, on ice. For even level reactions, the reaction master mix was prepared using 3.5 µl of dH_2_O, 0.5 µl of T4 DNA ligase buffer 10X, 0.5 µl of CutSmart buffer 10X (NEB cat. B7204S), 0.25 µl of T4 DNA ligase at 400 U µL^−1^ and 0.25 µl of SapI (NEB cat. R0569) at 10 U µL^−1^. Then, 5 µl of the reaction mix was combined with 5 µl of DNA mix for a reaction volume of 10 µl by pipetting, and incubated in a thermocycler as described previously (10). Next, 1 µl of the reaction mix was added to 50 µl of chemically competent TOP10 cells (ThermoFisher cat. C4040100;), incubated at 42°C for 30 s and left on ice for 5 min. A volume of 250 µl of Super Optimal broth with Catabolite repression (SOC) medium was added and cells were left incubating at 37°C for 1 h. Finally, cells were plated onto selective Lysogeny broth (LB)-agar plates supplemented with 1 mM Isopropyl b-_D_-1-thiogalactopyranoside (IPTG) (no. I6758; Sigma-Aldrich and 50 µg mL^−1^ of 5-bromo-4-chloro-3-indolyl-b-_D_-galactopyranoside (X-Gal) (no. B4252; Sigma-Aldrich).

### *E. coli* strain construction

Strain AKR1 was created using the GeneBridges “Quick & Easy Conditional Knockout Kit” following the manufacturer’s protocol. In order to make the linear DNA for chromosomal insertion, the FRT-PGK-gb3-hyg-FRT cassette, containing hygromycin resistance, was cloned adjacent to a constitutively expressed mRFP1, and the resulting DNA was further flanked by insulating terminators. The final insertion cassette was amplified by PCR with 75 bp DNA primers containing 50 bp ends designed to be homologous to a chromosomal region within the *arsB* locus, a previously characterised insertion site (24).

For the chromosomal insertion, *E. coli* TOP10 cells were transformed with plasmid pRedET, which harboured inducible Lambda Red machinery. The cells were induced and electroporated with purified DNA containing the insertion cassette. After recovery, transformants were plated onto LB plates containing 100 µg mL^−1^ hygromycin (Duchefa Biochemie). Chromosomal insertion was verified by Sanger sequencing of linear DNA amplified from 200 bp up- and down-stream of the insertion site (Source Biosciences). Hygromycin resistance was subsequently removed with the pCl-FLPe plasmid to produce the final strain AKR1, which showed constitutive expression of mRFP1. Correct antibiotic resistance removal was verified by Sanger sequencing of the genomic region.

### Productivity of assembly

Productivity of assembly was obtained as the number of resulting colonies from transforming 1 uL of reaction into 50 µL of AKR1 (*E. coli* TOP10-derived) homemade chemically competent cells with a competence of 2.5×10^8^ CFU µg^−1^ of pUC19. After heat-shock, 250 µL of SOC media was added to each transformation and incubated for 1 h at 37°C in a shaking incubator at 200 RPM. A volume of 10 µL of recovered cells was plated into LB plates supplemented with antibiotics, X-Gal and IPTG, and productivity of assembly was obtained by multiplying the number of colonies in the plate by 30.

### *E. coli* flow cytometry

Cell were inoculated overnight in M9 with antibiotics. A 1:50 dilution into fresh M9 media with antibiotics was performed in 96 well-plates and samples were left incubating for 3 h at 37 °C in a shaking incubator at 200 RPM. Samples were then measured in a BD FACS Aria II device (BD Biosciences, San Jose, USA) using a blue laser (ex. 488 nm) and a 530/30 nm filter for sfGFP and a 616/23 nm filter for mRFP1, recording 100,000 events for each sample.

### *E. coli* time-course plate fluorometry

Cell cultures of each plasmid transformed into AKR1 cells were grown for 8 h in M9-glucose media with kanamycin, spectinomycin or chloramphenicol antibiotics according to the resistance cassette of each plasmid. The cultures were diluted in 1:1000 in M9-glucose medium and 6 wells of Nunc Flat bottom 96-well black plates were loaded with 200 µl for each one. In the same way, 6 wells were loaded with AKR1 (GFP background), M9-glucose (OD600 background) and the plasmid pCAL1-sfGFP transformed into TOP10 cells (RFP background). Three experimental replicates were performed on different days (i.e. 18 wells per strain: 3 days with 6 technical replicates).

Plates were incubated at 37 °C in a Synergy HTX Plate Reader with continuous orbital shaking (282 cycles per minute) to perform the growth kinetics assays. Measurements of OD600 and fluorescence (RFP ex: 585 em: 620 / GFP ex: 485 em: 516) were made every 10 min for 24 h.

The obtained data was analysed using Python routines (see ‘Plate fluorometry data analysis’ on Supplementary Data) to obtain relevant parameters such as fluorescent expression and growth rate.

### Large-scale DNA assembly and CHEF gel

Four L3 fragments (L3-1_allx4 L3-2_allx4, L3-3_allx4, L3-4_sfGFP) were assembled in the presence of SapI and T4 DNA ligase under multiple conditions. A control reaction excluded T4 ligase to test for oligomerization. Other reaction conditions tested included all 4 fragments with full SapI/T4 DNA ligase mix, with variations in numbers of 16 °C/37 °C cycles (25x vs. 50x) and differing amounts of T4 DNA ligase (1x, 10 U µL^−1^ or 2x, 20 U µL^−1^). A reaction 1x master mix (0.5 µL T4 DNA ligase buffer 10x, 0.5 µL CutSmart buffer 10x, 3.25 µL ddH20, 0.5 µL of SapI at 10 U µL^−1^, and 0.25 µL or 0.5 µL of T4 DNA ligase at 400 U µL^−1^) was combined with a mixture of 1 µL ddH20 and 1 µL of each L3 plasmid for a total volume of 10 µL. Pulsed-field electrophoresis was carried out using a 1% agarose gel in 0.5x TBE buffer on the CHEF-DR III system (Bio-Rad). The entire 10 µL of each above described reaction was mixed with 10 µL of molten (55°C) 1% low-melting point agarose (Thermo-Fisher) and quickly pipetted into the appropriate lane after the gel was placed in the CHEF-DR III system and submerged in excess 0.5x TBE. One mm slices of the Midrange PFG markers (NEB cat. N0342S) were placed into the flanking lanes as size standards. Electrophoresis was carried out with the following parameters: initial switch time 1 s, final switch time 12 s, run time 12 h, 6 V cm^−1^, and a 120° angle. After electrophoresis, the gel was stained with a 0.5 µg mL^−1^ ethidium bromide solution in ddH20 for 30 min before imaging.

### Diatom transformation

Sub-strains CCMP632 and RCC2967 of *P. tricornutum* were obtained respectively from the National Center for Marine Algae and Protozoa (Bigelow Laboratories, Maine, USA) and Roscoff Culture Collection (Station Biologique de Roscoff, CNRS, Roscoff, France) and maintained in L1 medium prepared according to the NCMA recipe (https://ncma.bigelow.org).

Conjugation of episomes testing *de novo* assembly methods into *P. tricornutum* were carried out using the multi-well method described in Diner, et al. 2016 (25). Briefly, chemically-competent *E coli* DH5a cells harboring the pTA-MOB conjugation plasmid (26) were transformed with 10 ng of L2 episomes by heat-shock at 42°C, then plating outgrowths on LB medium containing 50 µg mL^−1^ kanamycin and 20 µg mL^−1^ gentamicin. Plates were incubated overnight at 37°C. Colonies from each plate were picked the next day and grown at 37°C overnight in liquid LB + 50 µg mL^−1^ kanamycin and 20 µg mL^−1^ gentamicin. These outgrowths were used to make glycerol stocks for storage at −80°C. Four days prior to conjugation, wild-type *P. tricornutum* cells were plated on 12-well tissue culture plates (Corning) filled with 3.5 mL of ½ L1 + 5% LB agar (250 µl of 1×10^8^ cells mL^−1^ cell resuspension, final cell count of 2.5×10^7^ cells/well) and incubated for 96 h on a 14:8 diel cycle at 22°C with a light intensity of approximately 50 µmol photons m^2^ s^−1^. The day before the conjugation, tubes containing LB + 50 µg mL^−1^ kanamycin and 20 µg mL^−1^ gentamicin were inoculated with scrapings from the glycerol stock tubes and allowed to grow overnight at 37°C. The conjugation, incubation, and selection plating on ½ L1 + 20 µg mL^−1^ phleomycin (InvivoGen) or 50 µg mL^−1^ zeocin were carried out exactly as described in Diner, et al. 2016 (25). Exconjugants were observed on selective plates after 10 days incubation at 22°C or 20 days at 20°C and picked directly into liquid L1 medium supplemented with 20 µg mL^−1^ phleomycin or 50 µg mL^−1^ zeocin for outgrowth of diatom colonies.

Episomes for testing the stability and repeatability of expression of uLoop plasmids (Section 6) were assembled with uLoop parts and transformed into PTA-MOB *E coli* via heat shock at 42°C as described above. Selection of colonies employed appropriate antibiotics (depending on receiver backbone used in assembly) and 20 µg mL^−1^ gentamicin and glycerol stocks were generated as described before. Conjugation into *Phaeodactylum* strains was carried out using the method described in Karas, et al. 2015 and plated on selective ½ L1 medium containing 50 µg mL^−1^ zeocin and 50 µg mL^−1^ ampicillin. Exconjugant colonies were visible on ½ L1-Zeo50 plates after 10-12 days incubation at 20°C under constant light.

### Transient expression in Arabidopsis mesophyll protoplasts

Plants were grown at 22°C in low-light (75 µmol m^−2^ s^−1^) and short-photoperiod (12 h: 12 h, light: dark) conditions. Well-expanded leaves from 4-week-old Arabidopsis plants (Columbia-0) were used for protoplast transfection. Isolation and PEG-mediated transformation (PEG 4000, Sigma-Aldrich) were made according to (27). For transfection, 6 µl of L2 plasmids (2 µg µl^−1^), isolated by a NucleoBond Xtra Midi/Maxi purification kit (Macherey-Nagel cat. 740410.50), were used. Transfected protoplasts were observed directly under epifluorescent microscopy after 12 h of light incubation in a Neubauer chamber (Hirschmann Laborgeräte, Eberstadt, Germany).

### Yeast transformation

*S. cerevisiae* cells were transformed following the lithium acetate/single-stranded carrier DNA/ polyethylene glycol method (28). The transformation was made according to the specifications for single plasmid by the addition of PEG 4000 50% (w/v), LiAC 1.0M, single-stranded carrier DNA (2.0 mg mL^−1^) and plasmid DNA plus sterile water (400 ng total DNA). Prior to the heat shock, cells were incubated at 30°C for 30 min. The heat shock was performed at 42°C for 30 min. Next, cells were plated on CSM-URA plates and incubated at 30°C for 2-3 days.

### Epifluorescence microscopy

Transfected protoplasts were examined using a Nikon Ni microscope (Minato, Tokyo, Japan) equipped with the following filter cubes: 49021 ET – EBFP2/Coumarin/Attenuated DAPI (excitation, 405/20 nm; dichroic, 425 nm; emission, 460/50 nm), 96223 AT-ECFP/C (excitation, 435/20 nm; dichroic, 455 nm; emission, 480/40 nm), 96227 AT-EYFP (excitation, 495/20 nm; dichroic, 515 nm; emission, 540/30 nm), and 96312 G-2E/C (excitation, 540/20 nm; dichroic, 565 nm; emission, 620/60 nm). Brightfield images of yeast cells were converted to greyscale for clarity (please refer to https://osf.io/kw4fh/ for all original files). All fluorescent images remained unedited.

### Confocal microscopy

Multispectral fluorescence microscopy was performed on a Leica TCS SP5 confocal microscope. mTagBFP2 was excited with a 405 nm laser, mTurquoise2 was excited with a 458 nm laser and Venus was excited with a 514 nm laser, collecting fluorescence in sequential mode using appropriate emission windows for each fluorophore (mTagBFP2, 450-470 nm; mTurquoise2, 470-490 nm; Venus, 520-540 nm). Images were then loaded into ImageJ and composed into a single multi-channel image 8-bit image. Contrast was auto-adjusted for each independent channel to span the range of the pixel intensity histogram.

### Flow cytometric analysis and cell sorting

Flow cytometric analysis of *P. tricornutum* grown in liquid culture was performed with an InFlux Cell Sorter (BD Biosciences, San Jose, CA, USA) equipped with 405 nm, 488 nm, and 640 nm lasers. Forward and side scatter and red fluorescence (692/40 nm filter) was excited by the 488 nm laser while blue fluorescence (460/50 nm filter) excited by the 405 nm laser. Sheath fluid was 3% NaCl, using an 85 µm nozzle, with sheath and sample pressures at 16 and 17 psi, respectively. Laser alignment was checked using 3 µm Ultra Rainbow fluorescent particles (Spherotech CAT# RCP-30-5A). Cytometer settings were maintained between WT and ex-conjugate *P. tricornutum* colonies analyzed on the same day. Between weeks, gains for fluorescence detectors were sometimes changed, but the relationship between signal and gain was determined using fluorescent particles and fluorescence was adjusted accordingly. Also, fluorescence of mTagBFP2-expressing exconjugates was always normalized with respect to ex-conjugates expressing the non-fluorescent protein LuxR and grown in the same conditions and run in parallel, to control for any potential variability in blue autofluorescence (from photosynthetic pigments, their degradation products, or other metabolites) related to episome maintenance and growth in antibiotics. In sorting experiments, 1 million blue fluorescent cells were sorted into 3 mL of L1 media+zeocin, and transferred to fresh medium approximately once a week.

## RESULTS

### uLoop system and library

uLoop was developed by decoupling plasmid propagation elements from the Loop assembly logic (**Figure 1A**). These propagation elements were converted and used as L0 parts during the uLoop assembly routine to create species-specific vectors from backbones optimised for efficient assembly (**Figure 1B**). Four uLoop vectors kits were constructed by implementing the Loop assembly logic (**Figure 1C**, **Supplementary Figure S1**) in derivatives of pCAMBIA, pJT170 (21), pSB4K5 (22) and pAN3945 (23). These vectors were modified by removing BsaI and SapI sites through a single nucleotide mutation of the recognition sequences. Vectors were minimised by removing sequences not related to plasmid replication and maintenance functions in *E. coli* (**Supplementary Figure S2**). Then, the Loop assembly schema was introduced into each vector to create four versions of uLoop vector kits (**Figure 1D**). uLoop vector kits were named according to the vector of provenance from which they were derived or their origin of replication: pCA (pCAMBIA-derived), pCO (ColE1 *ori*), pSB (pSB4K5-derived) and pAN (pAN3945-derived). Each kit version contains two sets of four vectors: four receiver plasmids of the odd levels (e.g.. pCAo-1, pCAo-2, pCAo-3, pCAo-4) and four receiver plasmids of the even levels (e.g. pCAe-1, pCAe-2, pCAe-3, pCAe-4). All odd levels of uLoop plasmids use kanamycin resistance for selection, and even levels use spectinomycin (pCA and pCO) or chloramphenicol resistance (pAN and pSB).

**Figure 1.**
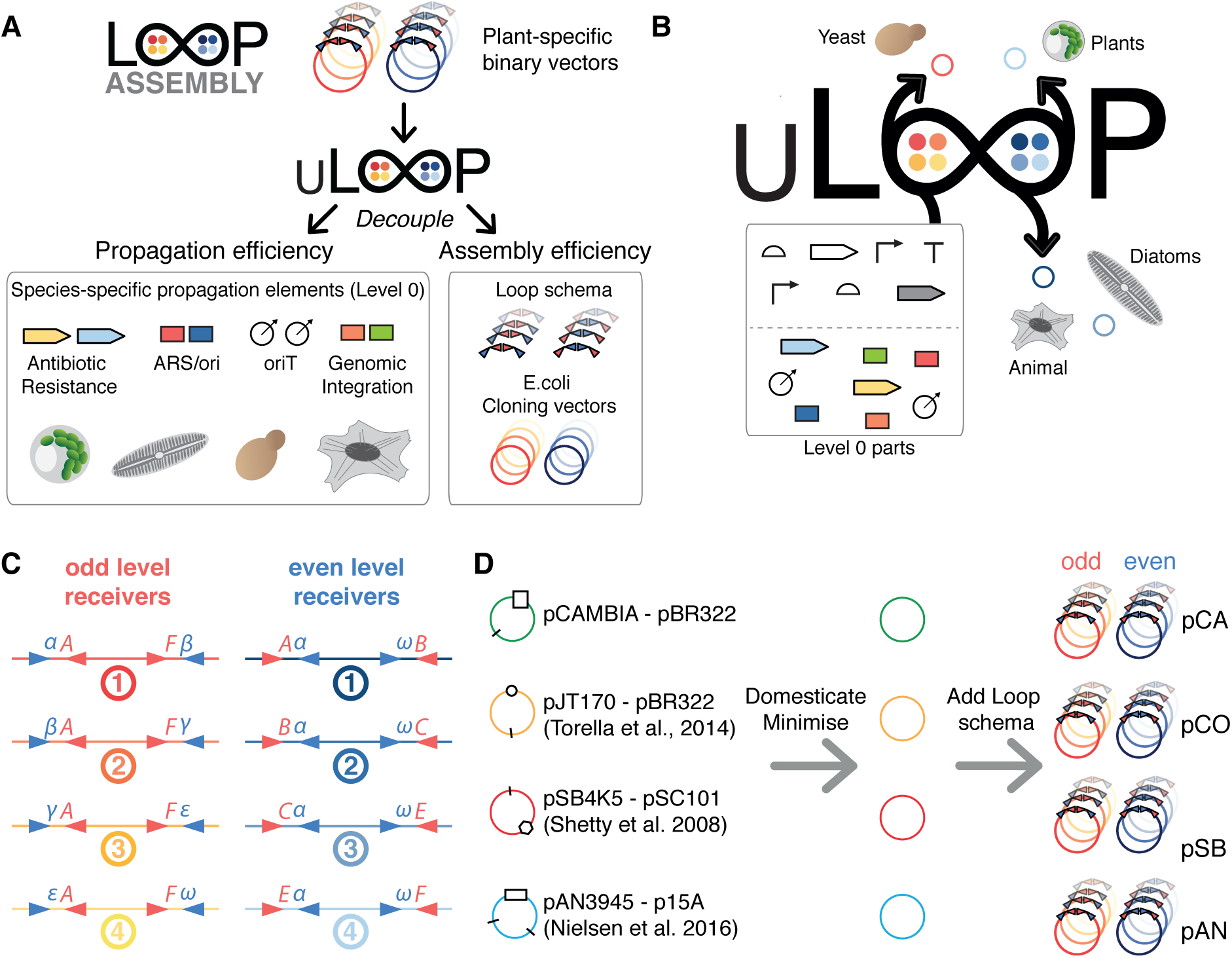
Universality of uLoop. (**A**) Decoupling of Loop assembly logic. uLoop decouples the propagation elements specific to the organism of choice from the universal assembly logic inbuilt in Loop vectors. (**B**) Creation of uLoop species-specific vectors from Loop. In uLoop, propagation elements required for transfer and maintenance of plasmids in target organisms are used as modular L0 parts. During the assembly routine, these cassettes are assembled into plasmids alongside with other transcription units to generate species-specific plasmids capable of transfer and propagation in the organism of interest. (**C**) Overhangs and restriction sites used in uLoop/Loop assembly. BsaI overhangs follow the common syntax (8,13), and SapI overhangs are the same as those described in Loop assembly (10), α(ATG), β(GCA), γ(TAC), ε(CAG) and ω(GGT). (**D**) Source of uLoop vector kits. Vectors from the synthetic biology community and the pCAMBIA vector were domesticated for BsaI and SapI and ‘minimised’ by removing elements not related to basic plasmid function. Two antibiotic resistance versions were generated and then the Loop schema was incorporated to generate odd and even plasmids for each version of vector kits.

The Loop assembly schema (10) was introduced to uLoop vectors with the only exception that in uLoop, all plasmids are flanked by the unique nucleotide sequences UNS1 (upstream) and UNSX (downstream) as described in Torella *et al*. 2014 (21), to enable PCR verification and sequencing of all plasmids with two standard oligonucleotides.

### Efficiency and productivity of assembly

Vector kits were tested for their capability to assemble DNA through the Loop assembly method. An assay using a sfGFP bacterial expression cassette was used to provide a visible phenotypic marker for tracking colonies likely to contain the assembled construct. The presence of lacZ expressing colonies (negative selection marker) would indicate undigested template while colonies not expressing sfGFP would report misassemblies. Based on this, we determined that efficiency of assembly, calculated as the percentage of sfGFP over total colonies not expressing the lacZ marker (white colonies), would provide a simple metric to compare the vector sets. We also counted the number of colonies present in each assembly, to compare productivity of assembly between the vector kits (**Supplementary Figure S3**).

Test assemblies were performed to characterise efficiency of assembly as the number of TUs per vector was increased. Assemblies were performed for a bacterial expression cassette of sfGFP in level 1 (L1-4_sfGFP), for a construct composed of three plant expression cassettes plus L1-4_sfGFP in level 2 (L2-4_sfGFP), and for a construct containing 12 plant expression cassettes plus the 4TU L2-4_sfGFP construct in level 3 (L3-4_sfGFP). Assemblies were performed into the fourth position of odd and even receivers of each kit (**Figure 2A**). For the L1-4_sfGFP construct, L0 parts were used (AB_J23101, BC_B0034m, CE_sfGFP and EF_B0015); for the L2-4_sfGFP construct, the L1-4_sfGFP plasmid was used along with three plant expression cassettes (pL1-1_35SmR3, pL1-2_35SmT2 and pL1-3_35SVe) used in our previous work for testing assembly efficiency (10); and the L3-4_sfGFP was assembled from three constructs containing four plant expression cassettes each (pL1-1_35SmR3, pL1-2_35SmT2, pL1-3_35SVe and pL1-4_35SmR3) and the L2-4_sfGFP construct (**Figure 2A**). After transformation, plates were scored for presence of colonies exhibiting blue coloration due to LacZ activity and fluorescence when imaged under UV light (**Supplementary Figure S3**).

**Figure 2.**
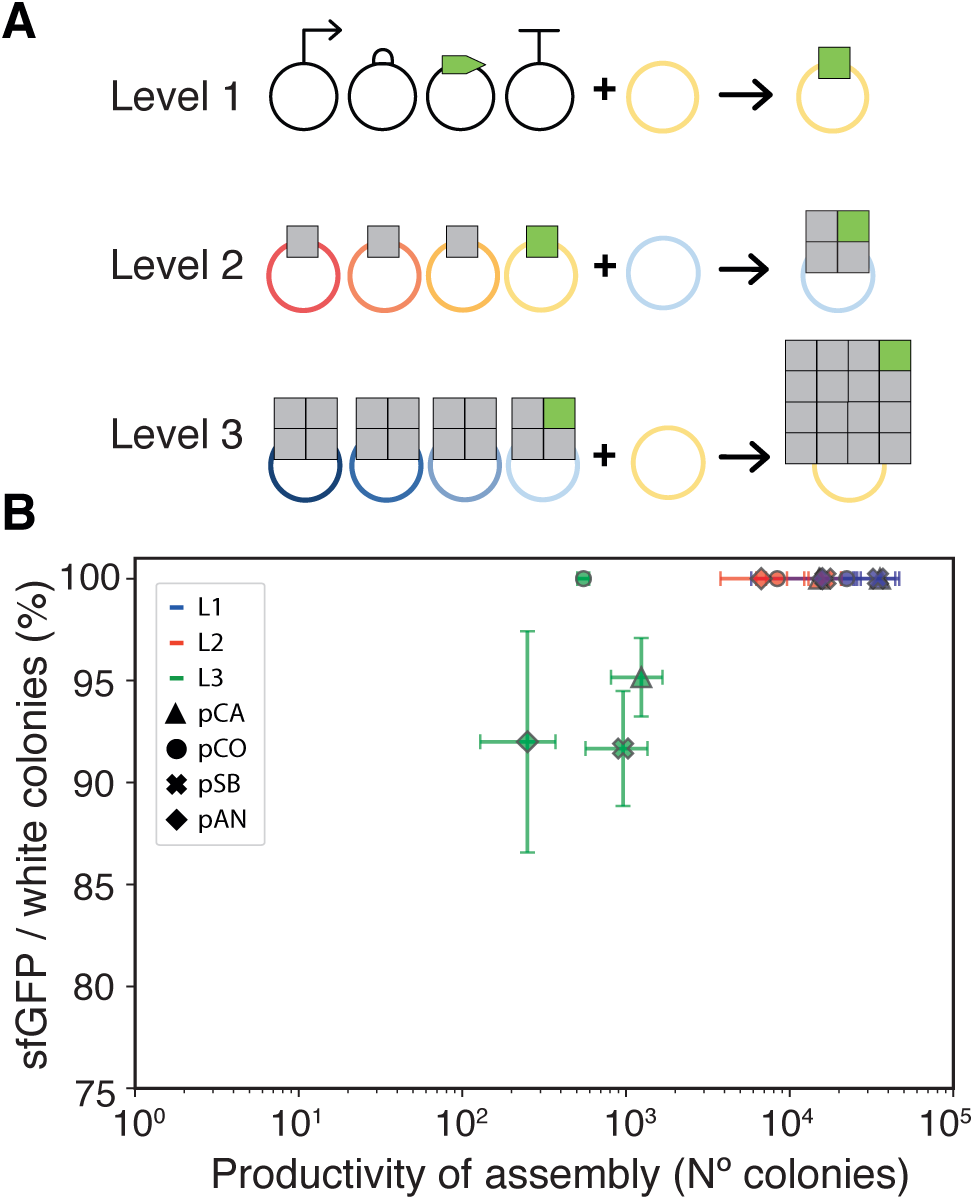
Efficiency and productivity of uLoop assembly. (A) Depiction of the assemblies tested for measuring efficiency and productivity of assembly. The Level 1 test assembly shows composition of L0 parts into a sfGFP TU in an odd receiver. The Level 2 test assembly shows the composition of 3 TUs with the L1_sfGFP TU into an even receiver. The Level 3 assembly shows the composition of 3 multi-TU constructs with the L2_sfGFP multi-TU construct into an odd receiver. (B) Efficiency versus productivity of assembly. Each vector kit is plotted using a different symbol, and levels of assembly are distinguished by color. Error bars represent 95% confidence intervals.

Levels L1 (3-4 kb) and L2 (9-10 kb) assemblies exhibited sfGFP expressing colonies predominantly over LacZ expressing colonies and no colonies lacking either sfGFP or lacZ expression were observed in any vector kit. For L3 (16 TUs, 32-33 kb), efficiency of assembly was 91% for pSB, 92% for pAN, 95% for pCA and 100% for pCO (**Figure 2B**). Greater differences were observed in the total number of non-negative colonies per plate where pSB and pCA vector kits showed substantially more colonies than pCO and pAN vectors (i.e. ∼2.2 fold more pSB colonies than pAN colonies in L1 and ∼2.5 fold in L2). Productivity in L3 was also reduced, averaging only 2.8% of the total number of colonies produced in L1 assemblies.

### Assembly integrity

Colonies showing a sfGFP phenotype were further tested to evaluate assembly integrity. Eight colonies per vector kit and level of assembly were analysed by means of restriction digest profiling. Resulting profiles from agarose gels were then scored against the expected restriction pattern for correct assemblies (**Table 1**, **Supplementary Figure S4A-C**). L1 assemblies showed 87.5% of expected profiles for pCA and pCO vectors, and 100% for pSB and pAN vectors. L2 assemblies showed 100% correct profiles for all vector kits. L3 assemblies evidenced a higher level of variability between vector kits, where the lowest percentage was exhibited by pCO and pSB (62.5%), followed by pCA (75%) and the highest percent by pAN (87.5%). L3 assemblies were repeated using BsaI-HFv2 and restriction profiles for sfGFP expressing colonies were assessed. Assembly integrity for BsaI-HFv2 did not seem to vary substantially from what was observed with BsaI, except for a slight improvement obtained for pCA, pCO and pAN (**Table 1**, **Supplementary Figure S4D**).

**Table 1.** Assembly integrity. * Integrity of assembly using BsaI-HFv2 instead of BsaI.

| Set | Level | Correct (n) | Incorrect (n) | Percentage (%) |
| --- | --- | --- | --- | --- |
| pCA | L1 | 7 | 1 | 87.5 |
|  | L2 | 8 | 0 | 100 |
|  | L3 | 6 | 2 | 75 |
|  | L3* | 7 | 1 | 87.5 |
| pCO | L1 | 7 | 1 | 87.5 |
|  | L2 | 8 | 0 | 100 |
|  | L3 | 5 | 3 | 62.5 |
|  | L3* | 5 | 3 | 62.5 |
| pSB | L1 | 8 | 0 | 100 |
|  | L2 | 8 | 0 | 100 |
|  | L3 | 5 | 3 | 62.5 |
|  | L3* | 4 | 4 | 50 |
| pAN | L1 | 8 | 0 | 100 |
|  | L2 | 8 | 0 | 100 |
|  | L3 | 7 | 1 | 87.5 |
|  | L3* | 8 | 0 | 100 |

We also tested different enzymes concentrations and reaction lengths for BsaI and SapI in an *in vitro* reaction assay that involved the assembly of 4 fragments. This was performed in the absence of a receiver vector to distinguish a full-length linear fragment by agarose gel electrophoresis. We used 0.25 U µL^−1^ or 0.5 U µL^−1^ of restriction endonuclease concentration in either a short cycle (1 min at 37 °C and 1.5 min at 16 °C) or a regular cycle (3 min at 37 °C and 4 min at 16 °C) for the 25 cycle reaction incubation. Resulting gels were analysed and the expected bands were quantified to determine formation of product in evaluated conditions. For BsaI, the condition that generated the highest amount of product was 0.5 U µL^−1^ of BsaI with the regular cycling conditions. For SapI, use of 0.5 U µL^−1^ increased the formation of product, but no discernible differences were obtained from the short and regular cycle (**Supplementary Figure S5**.

### Large-scale DNA assembly

*In vitro* reactions for large-scale DNA assembly were performed to assemble a 126 kb fragment (64 TUs) from four L3 parts, three containing 16 plant expression cassettes measuring approximately 32 kb each and L3-4_sfGFP. The reaction was performed using either 25 or 50 cycles of the assembly protocol (3 min at 37 °C and 4 min at 16 °C), varying the concentration of T4 DNA ligase to evaluate if this would enhance the formation of the intended product. Pulsed-field gel electrophoresis was conducted to visualise the reaction products, where it was possible to observe the 126 kb tetramer fragment in all reactions, including the monomeric, dimeric and trimeric conformations of the parts (**Figure 3**). Excess ligase did not seem to improve the reaction. This reaction was also conducted in the presence of a pCA even receiver vector and transformed into *E. coli*, however it was not possible to recover the intact plasmid from transformant colonies (data not shown).

**Figure 3.**
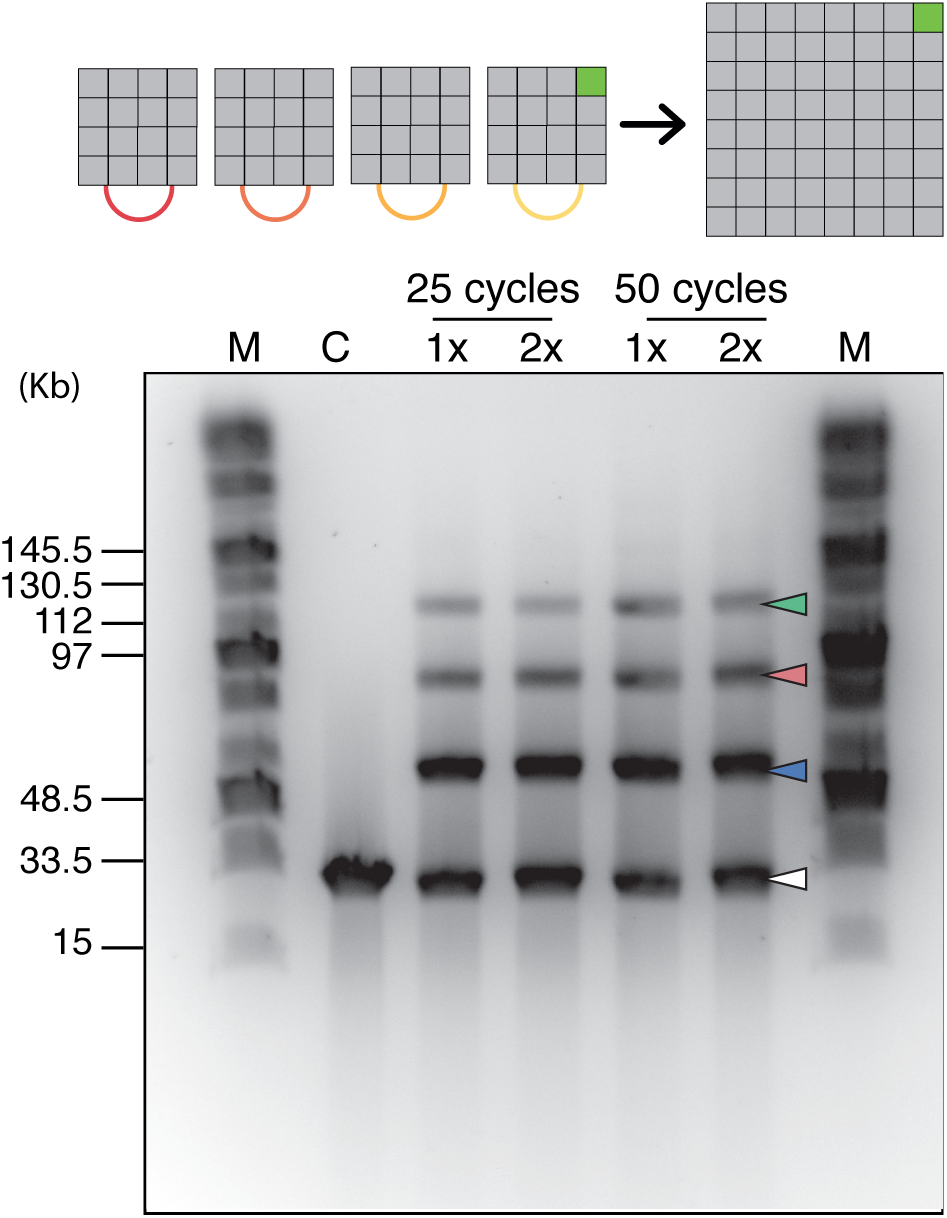
Large-scale DNA assembly. Four L3 parts were assembled in the absence of a receiver plasmid through a SapI-mediated Loop assembly reaction and products were analysed by pulsed-field gel electrophoresis. Lane headings: M, Midrange PFG marker. C, control reaction (L3 parts digested with SapI). Assembly reaction using 1x (10 U µL-1) or 2x (20 U µL-1) T4 DNA ligase. Image shown corresponds to an inverted photograph of the gel with adjusted contrast. The white arrow indicates the monomeric fragments, the blue arrow indicates the dimeric composites, the red arrow indicates the trimeric composites and the green arrow indicates the tetrameric full-length assembly.

### Performance of plasmids in *E. coli*

Performance of plasmids was evaluated in an AKR1 *E. coli*, a TOP10-derived strain harbouring a chromosomally integrated mRFP1 under constitutive expression, by means of time-course plate fluorometry and flow cytometry in mid-exponential phase. Plasmid-borne sfGFP expression was measured against mRFP1 expression as a proxy for plasmid versus genomic expression. L1, L2 and L3 levels for each vector set were contrasted to determine variability of expression in relation to plasmid size. Plasmid size varied from 3-4 kb in level 1 (only one sfGFP cassette) to 32-33 kb in level 3 (15 eukaryotic TUs and one sfGFP cassette), representing a ∼9 fold difference in plasmid size.

Plasmid-borne sfGFP to chromosomal mRFP1 ratio of expression measured by plate fluorometry and cytometry were consistent qualitatively (changes were in agreement in both assays). At a quantitative level, ratios varied according to detection parameters and thus cannot be directly compared. Although some of the vector kits showed significant variation between levels, these were not correlated with plasmid size (**Figure 4A** and **B**, **Supplementary Figure S6-S10** and **Supplementary Table S1**). Average sfGFP/mRFP1 fluorescence ratios for fluorometry and flow cytometry were highest for pSB vectors (4.51, 1.67), followed by pCA vectors (4.27, 1.50), then pCO vectors (2.37, 1.37) and finally pAN with the lowest average ratios (0.22, 0.57). A one-way ANOVA was conducted to compare the effect of plasmid size on plate fluorometry green-to-red fluorescent ratios of level L1, L2 and L3 (all vector kits grouped together). This analysis showed no statistically significant differences among L1, L2 and L3 (**Supplementary Table S2**). However, an ANOVA test comparing levels within each kit showed statistically significant differences across levels of the pCA kit (**Supplementary Table S3**). A Tukey’s honestly significant difference (HSD) post-hoc test showed that plate fluorometry green-to-red fluorescent ratios of level 3 were significantly lower than those of L2 and L1 for the pCA plasmid kit (**Supplementary Table S4**). Similar results were found for pCA comparing values obtained from flow cytometry experiments (**Supplementary Table S5** and **S6**). Additionally, similar analysis on flow cytometry data showed statistically significant higher ratios for level 2 of pAN with respect to level 1 and 2 of the same kit (**Supplementary Table S5** and **S6**, **Figure 4A** and **B**). We calculated the coefficient of variability (CV) for plate fluorometry and flow cytometry readings for each kit as a measure of internal variation. The lowest CV was found in pSB plasmids with a 15% and 21%, followed by pCO plasmids with a 23% and 19%, pCA plasmids with 36% and 41%, and finally pAN plasmids with 111% and 41%, for fluorometry and flow cytometry, respectively (**Supplementary Table S7**).

**Figure 4.**
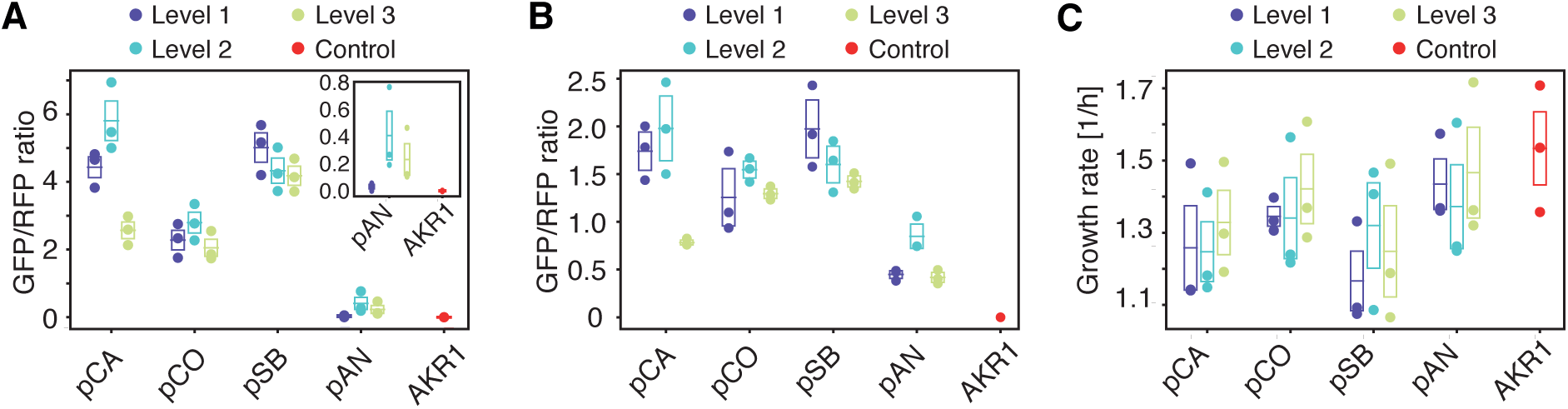
Performance of uLoop plasmids in *E. coli.* (**A**) Plate fluorometry of sfGFP/mRFP1 ratio of expression in *E. coli*. Performance of L1, L2 and L3 assemblies for all vector kits was assessed by measuring the ratio of expression of one copy of sfGFP per plasmid to a chromosomal mRFP1 cassette. These values were calculated from readings obtained over the full growth of cultures (see **Supplementary Information**). (**B**) Plasmid performance in *E. coli* measured by flow cytometry. Performance of L1, L2 and L3 assemblies for all vector kits was assessed by measuring the ratio of mean population values for green fluorescence (ex. 488 nm, em. 530 ± 15 nm) to red fluorescence (ex. 488 nm, em. 616 ± 11.5 nm) of cells expressing one copy of plasmid-borne sfGFP per plasmid and a chromosomal mRFP1 cassette. (**C**) Growth rates of L1, L2 and L3 assemblies for all vector kits and control AKR1 cells. Dots, boxes and line correspond to three measurements performed on different days; boxes and lines show standard error of the mean and mean, respectively.

Next, we analysed growth rates of bacterial cells carrying L1, L2 and L3 assemblies for all vector kits (**Figure 4C**, **Supplementary Figure S11-S15** and **Supplementary Table S8**). A Gompertz model was fitted to time-course densitometry measures to obtain growth rate parameters for each culture (see **Supplementary Information** for details). The growth rates of some plasmid cultures were lower than that of AKR1 control culture, however no statistically significant differences were detected among levels for each vector kit (ANOVA test, **Supplementary Table S9**).

### Performance of plasmids in *P. tricornutum*

To test the stability and repeatability of expression of uLoop plasmids in an eukaryotic background, a plasmid with mTagBFP2 under the control of histone H4 (pH4) promoter in the pSB vector, including oriT, the yeast centromere CEN-ARS (which allows plasmid maintenance in diatoms, (19)), and the pleomycin-resistance gene *shBle* (pSBL2-1_Pt-B). Separate conjugations were performed on *P. tricornutum* sub-strains CCMP632 and RCC2967 (both originating from strain CCAP1055/1). Eleven ex-conjugate colonies (six from CCMP632 and five from RCC2967) were tested from each conjugation and found to retain blue fluorescence for over four months while maintained in liquid culture. At all times tested, ex-conjugates displayed multi-modal histograms of blue fluorescence, indicating different expression phenotypes co-occurred in culture. The lowest mode had blue fluorescence that was indistinguishable from wild type *P. tricornutum* and ex-conjugates containing plasmids without mTagBFP2, while the highest modes showed over 50x greater blue fluorescence (**Supplementary Figure S16)**. Interestingly, the proportion of blue-fluorescence cells increased over time in four of the six ex-conjugates where less than 50% of cells showed blue fluorescence in the initial test (**Table 2**). Moreover, when high blue-fluorescent cells were separated by cell sorting and cultured, they retained the high blue-fluorescence phenotype when re-tested up to six weeks later (**Supplementary Figure S17**, **Table 2**).

**Table 2.**
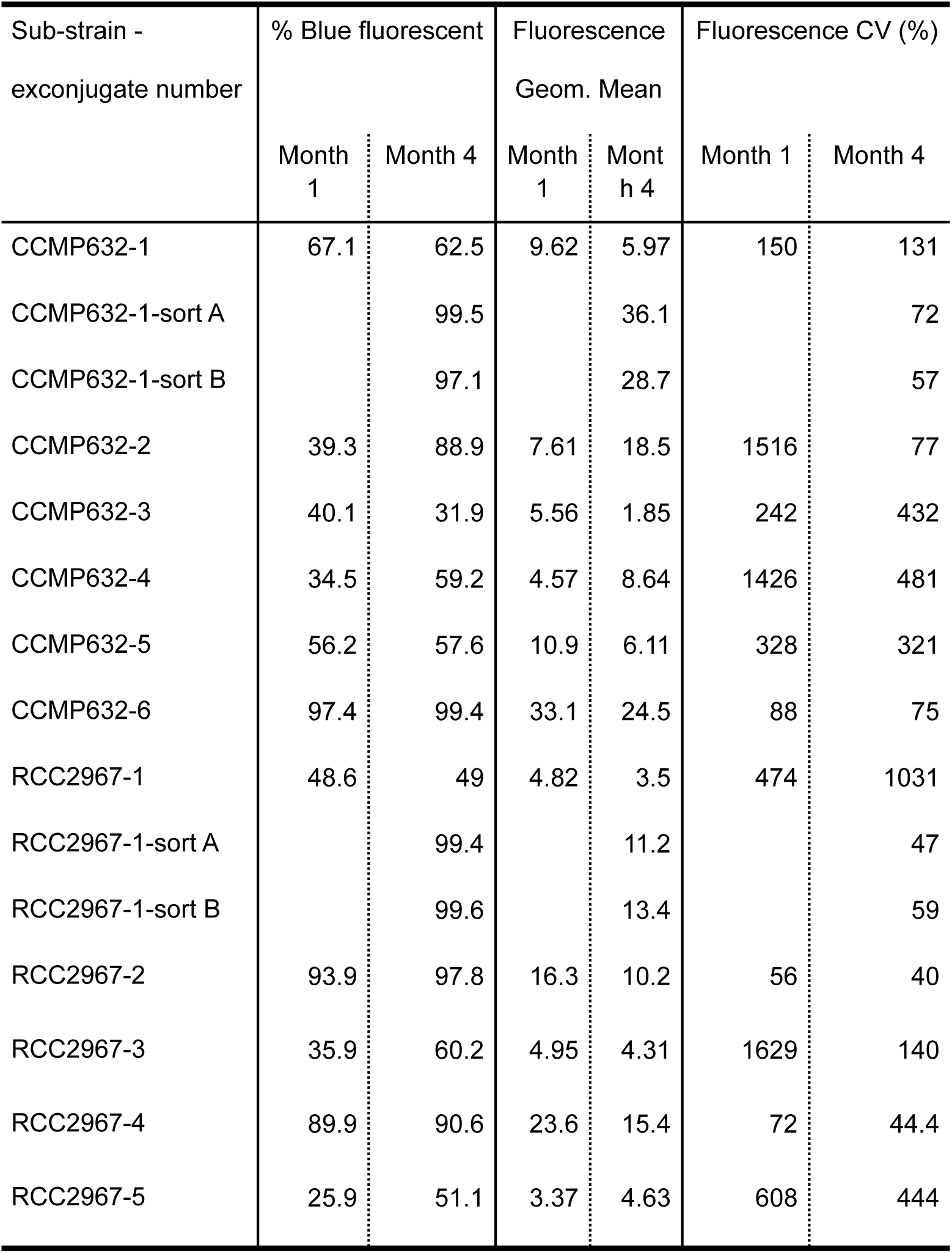
Stability of uLoop plasmid in *P. tricornutum*. Fluorescent phenotypes of 11 pSBL2-1_Pt-B ex-conjugate colonies from conjugations of two separate sub-strains (CCMP632 and RCC2967) were tested 3 months apart. The highest sub-populations of blue fluorescent cells were sorted on two independent days at month 2.5 (sort A and sort B) from one exconjugate colony of each conjugation and re-tested at month 4 (see **Supplementary Figure S17**). Reported are the % of cells showing blue fluorescence above the 95th percentile for control (non-BFP) exconjugates, the geometric mean of blue fluorescence (normalised to control exconjugates), and the coefficient of variation (CV) of blue fluorescence within each exconjugate. Fluorescence was measured logarithmically.

| Sub-strain -<br>exconjugate number | % Blue fluorescent |  | Fluorescence<br>Geom. Mean |  | Fluorescence CV (%) |  |
| --- | --- | --- | --- | --- | --- | --- |
|  | Month 1 | Month 4 | Month 1 | Month 4 | Month 1 | Month 4 |
| CCMP632-1 | 67.1 | 62.5 | 9.62 | 5.97 | 150 | 131 |
| CCMP632-1-sort A |  | 99.5 |  | 36.1 |  | 72 |
| CCMP632-1-sort B |  | 97.1 |  | 28.7 |  | 57 |
| CCMP632-2 | 39.3 | 88.9 | 7.61 | 18.5 | 1516 | 77 |
| CCMP632-3 | 40.1 | 31.9 | 5.56 | 1.85 | 242 | 432 |
| CCMP632-4 | 34.5 | 59.2 | 4.57 | 8.64 | 1426 | 481 |
| CCMP632-5 | 56.2 | 57.6 | 10.9 | 6.11 | 328 | 321 |
| CCMP632-6 | 97.4 | 99.4 | 33.1 | 24.5 | 88 | 75 |
| RCC2967-1 | 48.6 | 49 | 4.82 | 3.5 | 474 | 1031 |
| RCC2967-1-sort A |  | 99.4 |  | 11.2 |  | 47 |
| RCC2967-1-sort B |  | 99.6 |  | 13.4 |  | 59 |
| RCC2967-2 | 93.9 | 97.8 | 16.3 | 10.2 | 56 | 40 |
| RCC2967-3 | 35.9 | 60.2 | 4.95 | 4.31 | 1629 | 140 |
| RCC2967-4 | 89.9 | 90.6 | 23.6 | 15.4 | 72 | 44.4 |
| RCC2967-5 | 25.9 | 51.1 | 3.37 | 4.63 | 608 | 444 |

### Library of parts and nomenclature

A core library of parts was generated to enable the use of uLoop plasmids in different organisms. The library was generated using the common syntax for standardised DNA components (13). Parts domesticated for this work include promoters, coding sequences and terminators for the composition of TUs for bacteria, yeast, *P. tricornutum* and *A. thaliana*. Other parts not functionally described as standardised components (e.g. origins of transfer, centromeres, TUs encoding resistance genes for selection) were domesticated using common syntax overhangs for their composition. L0 parts, as well as L1 and L2 assemblies, used in this work along with metainformation are listed in **Supplementary Table S10**, **S11 and S12**, respectively. Furthermore, a virtual repository is currently under development to access information and parts’ documentation in www.uloop.org.

To denominate L0 parts and their composability we established a simple nomenclature describing overhangs flanking each part. Parts’ names are preceded by letters to depict the overhangs that flank them (e.g. AC_CEN-ARS-HIS, CE_oriT, EF_PtBle), being A & F terminal overhangs. We used a constrained implementation of the common syntax at the part level to limit redundancy in the library. For promoters, the A and C overhangs were used, while A and B overhangs were used when a N-terminal fusion was required. B and C overhangs were used for N-terminal tags. For CDSs, C and D overhangs were used. The STOP codon (if present) was removed and GC nucleotides were included to encode alanine and glycine with the D overhang. For C-terminal tags, D and E overhangs were used except when no C-terminal tag was required, in which case a DE_3xSTOP codon part was used. For 3’UTRs and terminators, E and F overhangs were used.

### Use of uLoop vectors across biological kingdoms

To demonstrate the universality of uLoop across different kingdoms, vectors containing transformation/transfection/conjugation elements were generated to allow transfer of DNA into different target organisms. Elements used in current transformation methods for each specific target organism were included in the vectors during the assembly routine along with TUs encoding fluorescent proteins. Then, vectors were transferred to the target organisms using described protocols and their function was assessed by fluorescence microscopy. A pCAL2-1_FPrep vector for transformation into *P. tricornutum* was constructed by hierarchical assembly. A module containing the origin of transfer (oriT), the yeast centromere CEN-ARS-HIS and a bleomycin resistance cassette (*shBle*) was assembled along with two fluorescent proteins tagged with localisation signals and a cytoplasmic fluorescent reporter. The fluorescent reporters corresponded to a pH4-driven (29) mTurquoise2 fluorescent protein fused to a mitochondrial localisation tag (30), a pNR-driven (31) Venus fluorescent protein fused to a peroxisomal localisation tag (32) and a p49202-driven (V. Belinski and C. Dupont, unpublished results) cytoplasmic mTagBFP2 fluorescent protein. The native terminator corresponding to each promoter was used. The plasmid was transformed into an *E. coli* strain containing pTA-MOB and then conjugated into *P. tricornutum*, using 20 µg mL^−1^ phleomycin as selection. Ex-conjugant colonies were propagated into liquid culture and then imaged using laser-scanning confocal microscopy. Microscopy inspection showed expression of all fluorescent reporters where Venus fluorescence was localised in discrete puncta in the vicinity of the nucleus; mTurquoise2 was found in elongated structures across the cell and mTagBFP2 was localised throughout the cytoplasm (**Figure 5A**, **Supplementary Figure S18A**).

**Figure 5.**
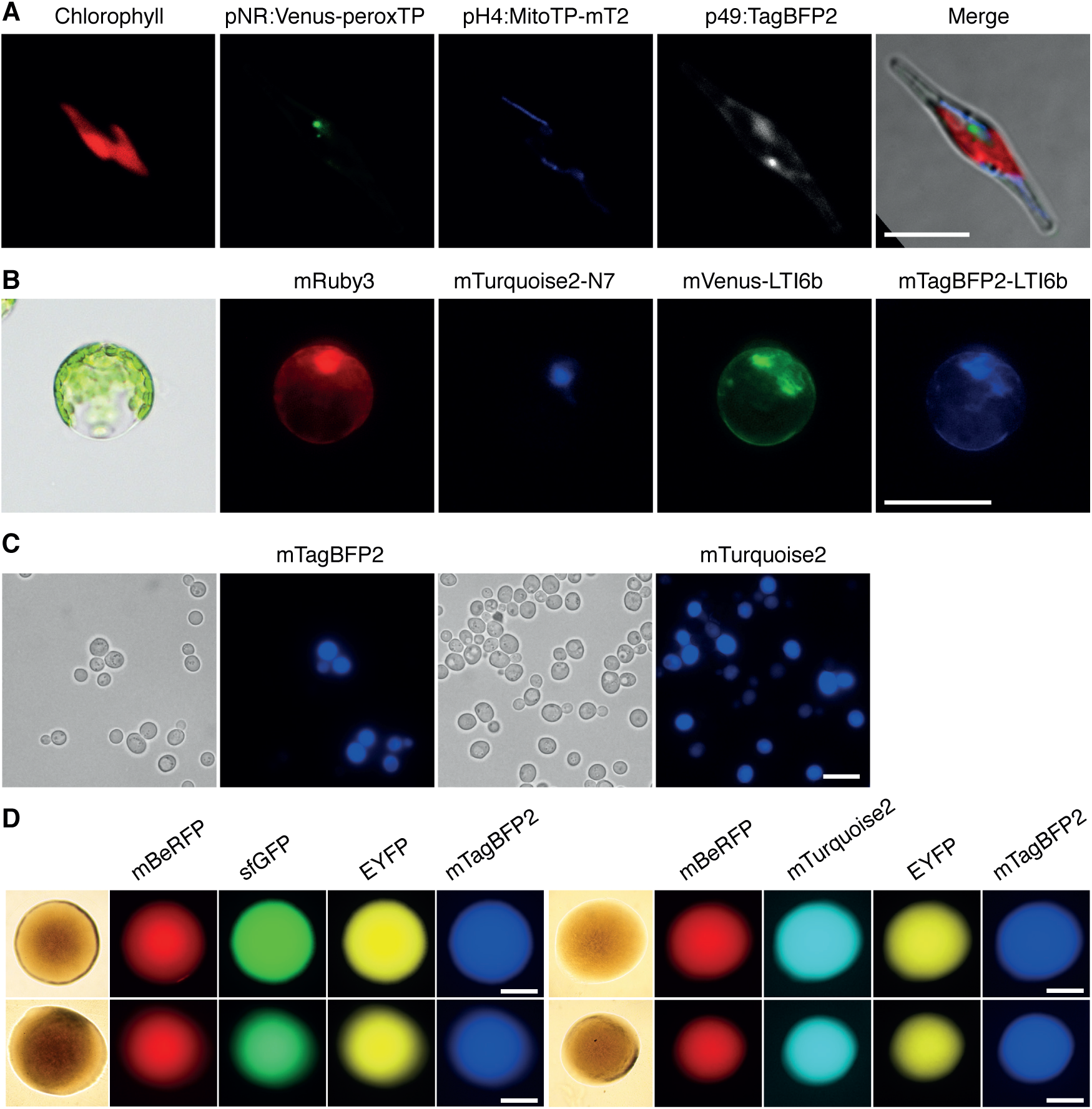
Use of uLoop vectors across multiple organisms for multi-spectral fluorescence. (**A**) Expression of three fluorescent reporters in *P. tricornutum.* From left to right, Chlorophyll fluorescence, mVenus fluorescent protein fused to a peroxisomal localisation tag, mTurquoise2 fluorescent protein fused to a mitochondrial localisation tag and mTagBFP2 fluorescent protein expressed in the cytoplasm. Scale bar = 10 µm. (**B**) Expression of four fluorescent reporters in protoplasts of *A. thaliana* from pCAL2-1_4xFP. From left to right, mRuby3 fluorescent protein expressed in the cytoplasm, mTurquoise2 fluorescent protein fused to the nuclear localisation tab N7, Venus fluorescent protein fused to plasma-membrane localisation signal LTi6b and mTagBFP2 fluorescent protein fused to plasma-membrane localisation signal LTi6b. Scale bar = 50 µm. (**C**) Expression of two fluorescent reporters in *S. cerevisiae.* Left, transmitted light microscopy and mTagBFP2 expression from pSB plasmid. Right, transmitted light microscopy and mTurquoise2 expression from pCA plasmid. Scale bar = 10 µm. (**D**) Expression of four fluorescent reporters in colonies of *E. coli*. Left: mBeRFP, sfGFP, EYFP and mTagBFP2 fluorescent protein expression from pCA (top) and pSB (bottom) uLoop vectors. Right: mBeRFP, mTurquoise2, EYFP and mTagBFP2 fluorescent protein expression from pCA (top) and pSB (bottom) uLoop vectors; Bottom: pSB. Scale bar = 500 µm.

To evaluate the functionality of a multi-TU construct in a plant organism, a transient gene expression system using Arabidopsis mesophyll protoplasts was used (27). Protoplasts were transfected with a construct encoding 4 constitutive fluorescent reporters in pCA. Reporters used correspond to a cytoplasmic mVenus, mTagBFP2 fluorescent protein fused to a Lti6b membrane localisation tag (33), a mTurquoise2 fused to a N7 nuclear localisation tag (33) and a cytoplasmic mRuby3. In all cases, fluorescent proteins were driven by the Arabidopsis UBQ10 promoter, which provides stable expression of fluorescent reporters in plants (34). Transfected protoplasts were examined through fluorescence microscopy where mRuby3 fluorescence was found in the cytoplasm, mTurquoise2 fluorescence was concentrated only in the nucleus, and mVenus and mTurquoise2 fluorescence had similar localisation patterns in the area surrounding the nucleus and in the membrane (**Figure 5B** and **Supplementary Figure S18B**).

Next, we tested uLoop vectors in yeast. Two vectors (pCAL2-1_Yeast-mT2 and pSBL2-1_Yeast-B) were constructed in pSB and pCA receivers to evaluate their functionality in *S. cerevisiae*. These vectors contained an origin of replication for yeast (2u or CEN), a selection marker (URA3) and a single fluorescent protein (mTurquoise2 or mTagBFP2). The expression of the fluorescent markers was confirmed by fluorescence microscopy (**Figure 5C**). Finally, we tested uLoop vectors in *E. coli* for multi-spectral fluorescent protein expression. We used two sets of four constitutively-driven fluorescent proteins: mBeRFP, EYFP, mTagBFP2, and either sfGFP or mTurquoise2 assembled on a pCA or a pSB vector. These four L2 constructs (pCAL2-1_RYBG, pCAL2-1_RYBmT2, pSBL2-1_RYBG and pSBL2-1_RYBmT2) containing different combinations of constitutively-driven fluorescent proteins were inspected by fluorescence microscopy, where expression of all fluorescent reporters was observed (**Figure 5D**, **Supplementary Figure S18C**).

## DISCUSSION

Here we describe the uLoop suite of vectors for universal DNA assembly. These vector kits, derived from plasmids widely used in synthetic biology, provide a repertoire of *ori* options for plasmid copy-number control in *E. coli*, as well as a modular mechanism for incorporating propagation elements for the customisation of vectors depending on the destination organism. The use of pCambia vectors was due to their freedom-to-operate policy while the use of pSB4K5 backbone was motivated by the prospect of providing resources to the international Genetic Engineering Machine (iGEM) community, which incentivises the advancement of Synthetic Biology through a worldwide student competition and open access to genetic resources. In this respect, the pSB vector kit could facilitate the exploration of new model organisms for iGEM projects (e.g. diatoms). Further, we have evaluated the capacity for assembly for each vector kit and showed that all are capable of reliably assembling constructs containing up to 16 TUs. All vector kits showed very high efficiency, productivity and integrity of assembly for plasmids up to 4 TUs, showing slight variation between the measures evaluated. Assembly of L3 constructs up to 16 TUs (54 DNA parts) showed a lower rate of efficiency, productivity and integrity of assembly, yet complete constructs were still easily obtained. Notably, even in the worst case, half of the putative positive colonies screened showed the correct assembly of 16 TUs. The decrease in the measures evaluated can be due to the size of the construct, since transformation efficiency decreases substantially past a certain size (35,36). At that point, the un-cut template has an advantage for transformation, influencing both the efficiency and the productivity of assembly.

To test uLoop’s capacity for large-scale DNA construction, assemblies were conducted up to a level 4 linear fragment measuring 126 kb (64 TUs). Although we evidenced the assembled full-length product by electrophoresis, it was not possible to obtain transformants harbouring the complete construct. This can be due to the high content of direct repeats of the assembly which could be lost *in vivo*, which could also explain the lesser integrity of assembly observed for L3 constructs. The L4 construct composed of 64 TUs contained 63 CaMV35S promoters, 31 mRuby3 CDSs, 16 mTurquoise2 CDSs, 16 Venus CDSs, 32 N7 tags and 63 nos terminators, aside from the sfGFP expression cassette. Direct-repeats are known to induce deletions in *E. coli* (37). Repeats could be lost during replication if the construct exerts a selection pressure for DNA replication. In a pBR322 origin containing-plasmid with an average copy-number of 20, the ∼128 kb plasmid (126 kb + backbone) would amount to roughly half of an *E. coli*’s genome in terms of DNA content. Presence of the complete fragment demonstrates that uLoop enables the construction of large DNA, however it remains unclear if absence of direct repeats could allow recovery of such DNA in *E. coli* due to the metabolic burden demanded by the construct size and copy-number of the vector used. Alternatively, a lower copy-number origin of replication (e.g. pDestBAC, (21)) could lead to higher stability enabling such DNA to be replicated. Nevertheless, the assembly of constructs harbouring up to 16 TUs (L3 constructs) would largely cover most current cloning needs for both genetic engineering and synthetic biology.

Characterisation of plasmid performance in bacteria showed that the effects of construct size on sfGFP expression levels can vary for each plasmid kit. In our experiments, variability was higher between vector kits than between plasmid levels. Further, pSB and pCO kits showed a low level of variability of expression between levels according to coefficients of variation, suggesting that these vector kits can provide more consistent expression in *E. coli* regardless of their size.

uLoop plasmids performed efficiently in each eukaryote system tested, demonstrating the versatility and broad applicability of the system. The target organisms were selected to represent three eukaryotic superkingdoms which diverged at the base of the eukaryotic tree of life (38), with the yeast *S. cerevisiae* from the Opisthokonts (animals and fungi), *A. thaliana* from the Archaeplastida (plants, green algae, red algae) and the diatom *P. tricornutum* from the SAR (Stramenopile, Alveolate, Rhizaria) superkingdom. In all these organisms, multispectral fluorescence imaging showed that multiple TUs are successfully expressed from uLoop plasmids. Special attention was devoted to testing uLoop in *P. tricornutum*, recognising the special needs for genetic tools in such emerging model organisms and the need for new marine microbial model systems (39). The uLoop plasmids were efficiently introduced in *P. tricornutum*, in which exconjugants exhibited variable levels of fluorescent protein expression. Nevertheless, high-level expression phenotypes were maintained and even increased in relative abundance with time and, furthermore, the high-expression phenotype was very stable after cell sorting. This high stability over time is consistent with their design to function as episomes in *P. tricornutum* and suggests that they do not present an excessive metabolic burden to cells.

A cross-kingdom DNA assembly method has also been described recently (40) but, unlike uLoop, this system is limited to 5-part vector assemblies. Any extension of the number of TUs assembled with this system appears to translate into a one-to-one increase in the number of LII plasmids required, as well as modifications of the LIII plasmids for each extension intended. This linear scaling of vectors needed per TU is present across all the GG-based methods described so far, apart from Loop (10). The recursive nature of the Loop schema used in uLoop solves this problem by inverting the recognition sites of BsaI and SapI in each level, which permits reusing the same vector sets for multilevel assemblies. This feature also provides a generalised framework for universal parts to be used in any position. For instance, a unique universal spacer linker can be used in any of the four position, and transcription factor binding elements can be used to create level 0 combinatorial promoter libraries (10). Moreover, Chiasson et al. (40) described several vector backbones with different propagation elements for specific hosts, which indicates that the number of plasmids would further increase if new target species are to be used. In contrast, uLoop decouples the two processes, delivering species-agnostic plasmids optimised for assembly in *E. coli* that can be customised for target hosts through basic (level 0) components. In addition to reducing the number of vectors, this design avoids the need for creating *de novo* backbones and re-characterising their assembly efficiency. Thus, while most GG-based methods offer an overwhelming number of plasmids for large assemblies specific for different hosts, uLoop only uses the same two sets of four plasmids for any target organism. This significant reduction of complexity has proven critical for the rapid adoption of the system in several institutions and countries.

Together with the development of the uLoop vector kits, we generated a basic set of DNA parts (promoters, fluorescent proteins and terminators, among others) for this work. We expect uLoop to boost the growth of species-specific level 0 part collections for different hosts. To address the problem of traceability of DNA parts, we are developing a virtual repository (www.uloop.org) to collect information such as sequences, descriptions, submitter, source of DNA, lab of origin in order to help with accessibility of parts’ information and requests for materials. The establishment of digital tools and a crowdsourced repository seeks to promote a distributed growth of libraries of openly shared and well-documented parts for different organisms. Adoption of uLoop in different model systems will provide the repository with an influx of DNA parts from diverse phylogenetic origin (such as transcription factors and repressors) and other genetic tools specific to model organisms. These parts would be useful for synthetic biology and genetic circuit engineering, and their availability would encourage further exchange between fields of biology.

Finally, the work described here was conducted in several laboratories around the globe, where the simplicity of the method facilitated its rapid implementation and experimental work. A key element for rapid and wide distribution of the uLoop vector kits in the future is the Open Material Transfer Agreement, a legal tool that enables seamless sharing and use of biological materials (15). Open access to materials and freedom to operate will be crucial to maximise interoperability and enable the growth of distributed uLoop libraries and toolkits in different countries around the world.

## AVAILABILITY

Data analysis and plotting scripts are provided as Jupyter Notebooks in a GitHub repository. (https://github.com/Prosimio/Plate_reader_analysis).

DNA sequences are provided as Supplementary Files.

## SUPPLEMENTARY DATA

Supplementary Data is supplied as a PDF file.

## ACKNOWLEDGEMENTS

We would like to acknowledge Patrick Brunson for helping with *P. tricornutum* promoter and terminator sequences, Diego Bustos and Javiera López for advice and support on yeast transformation methods. We would like to thank Andrea Calixto for kindly proofreading the manuscript and providing useful comments.

## FUNDING

This work was funded by the Gordon and Betty Moore Foundation [GBMF5007.01 to CLD and VB, GBMF4981.01 to FF and PvD]. BP was supported by the Programa de Apoyo a Centros con Financiamiento Basal [grant number AFB 170004]. AK was supported by the National Institutes of Health [grant number 1R01DK11077001A1]. FF, BP, TM and IN were supported by Instituto Milenio iBio and PvD by the Instituto Milenio de Oceanografía [grant number IC120019] of the Iniciativa Científica Milenio MINECON. IN was also supported by a PUC VRI PhD studentship and AC was supported by CONICYT scholarship for doctoral degrees in Chile [grant number 21150778].

## CONFLICT OF INTEREST

None.

## Notes

#### Summary of Updates

Supplementary information ammended

https://osf.io/kw4fh/

