## Supplementary information for "Universal Loop assembly (uLoop): open, efficient, and species-agnostic DNA fabrication"

### Supplementary Figures

### Supplementary Tables

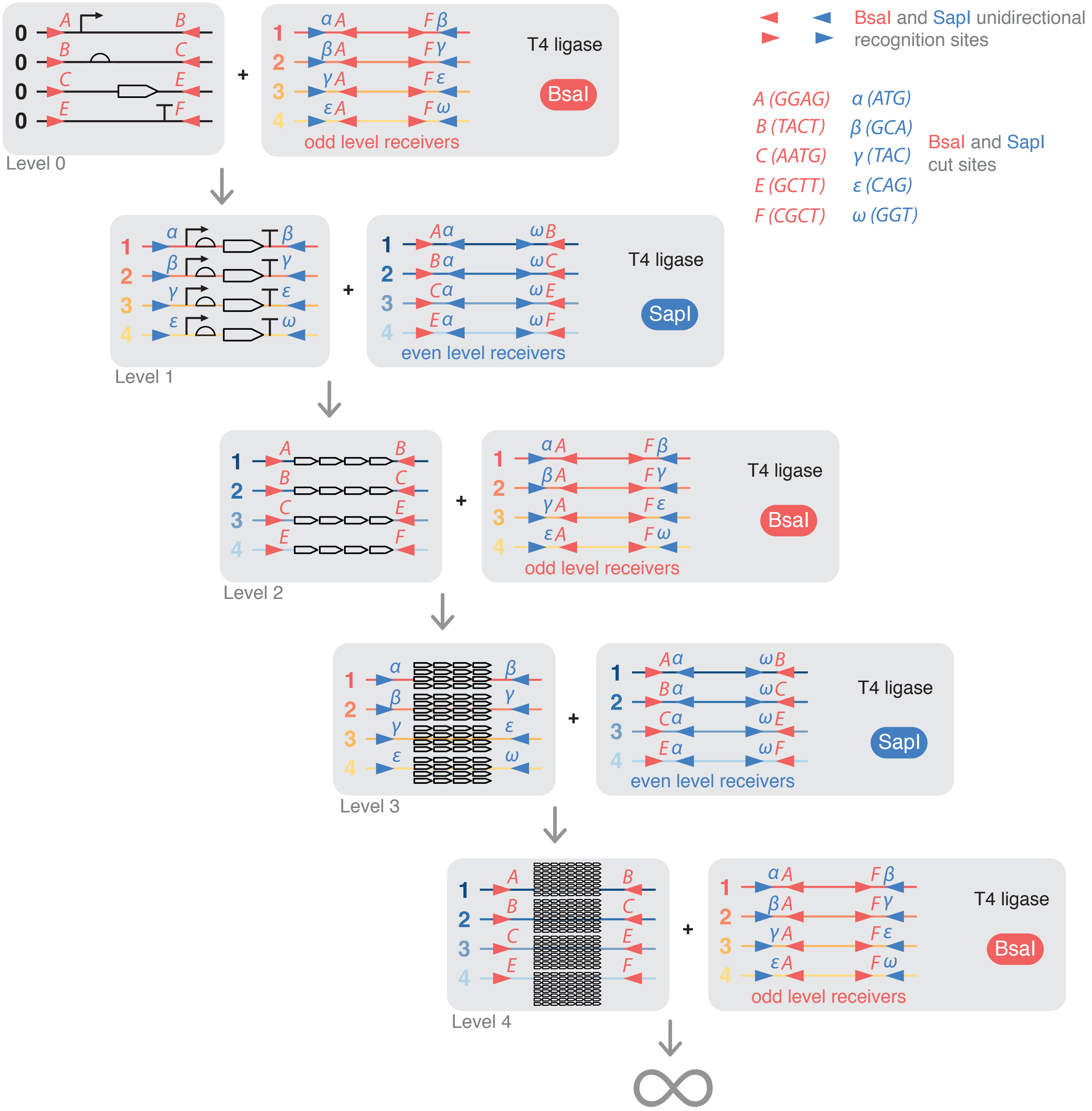

**Supplementary Figure S1. Diagram of uLoop assembly logic.**

uLoop is composed of two sets of four plasmids, called even and odd receivers (shown in red and blue, respectively). Vectors of exponentially increasing size are built by alternating BsaI and SapI Golden Gate assembly reactions through the repeated use of odd and even vectors. The use of inverted orientations for BsaI and SapI recognition sites in odd and even receivers permits the product of one reaction to become the substrate for the next reaction at the following level. Up to four assemblies can be conducted in parallel per level, allowing DNA elements to be composed in an exponential manner of 4 genetic modules per assembly round. BsaI overhangs follow the common syntax [1, 2], and SapI overhangs are the same as those described in Loop assembly [3], $\alpha(ATG), \beta(GCA), \gamma(TAC), \alpha(CAG)$ and $\omega(GGT)$ for hierarchical and sequential assembly in repetitive loops.

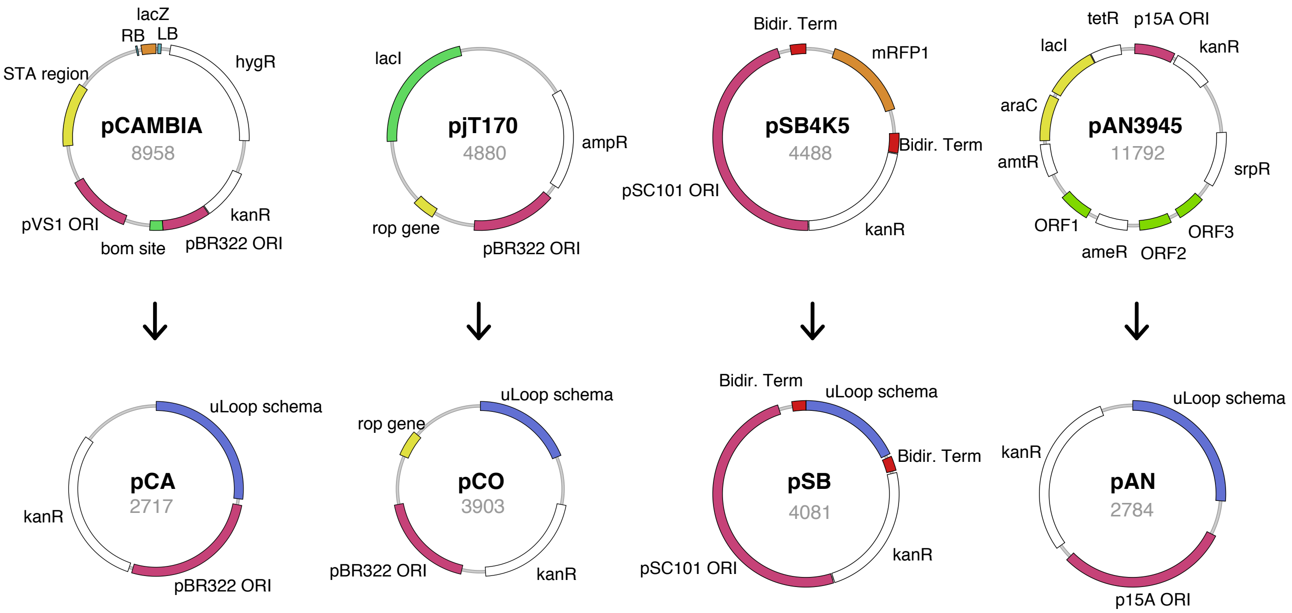

**Supplementary Figure S2. Vector backbone refactoring.**
Vectors from the synthetic biology community (pjT170, pSB4K5 and pAN3945) and the pCAMBIA vector were domesticated for BsaI and SapI and 'minimised' by removing elements not related to basic plasmid function. Numbers below the vector name denote the size in base pairs.

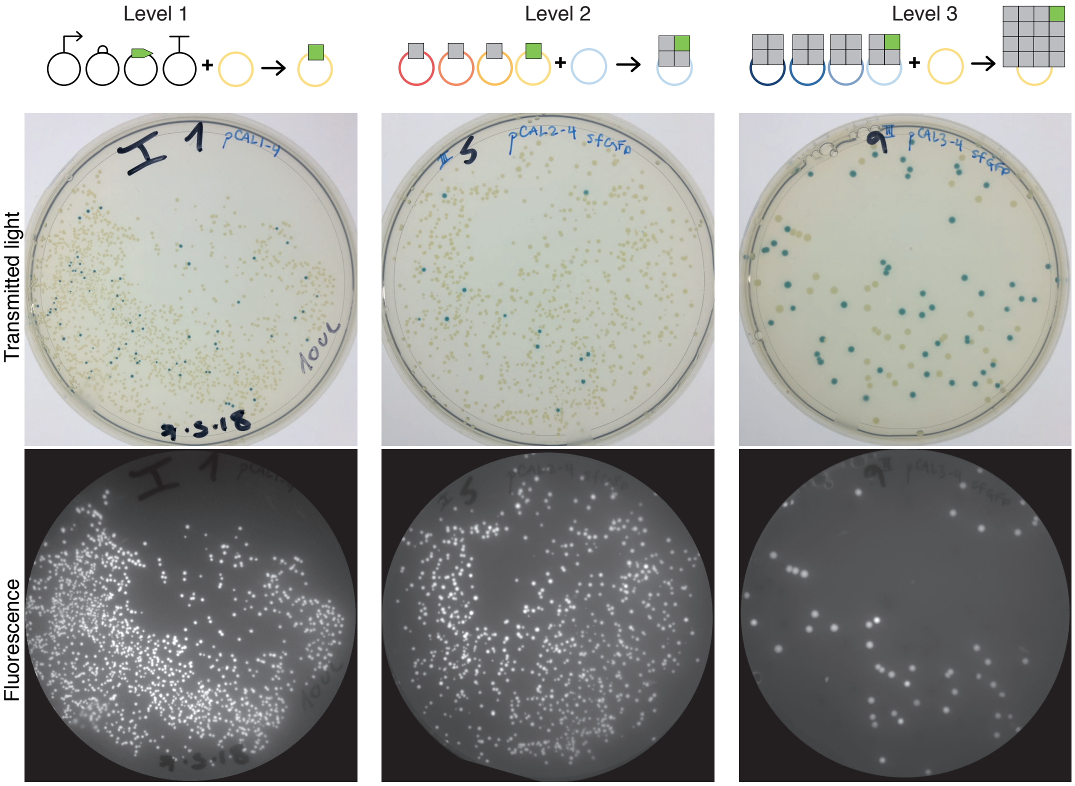

**Supplementary Figure S3. Plate images for efficiency and productivity of assembly.**
Resulting plates for L1 (left), L2 (middle) and L3 (right) assembly reactions are shown. Transmitted light images show the presence of blue colonies that correspond to the negative background of the reactions given by the presence of the negative selection marker LacZ (top panels). In each assembly a single bacterial sfGFP expression cassette enables visualisation of fluorescent colonies in plates under UV illumination (bottom panels).

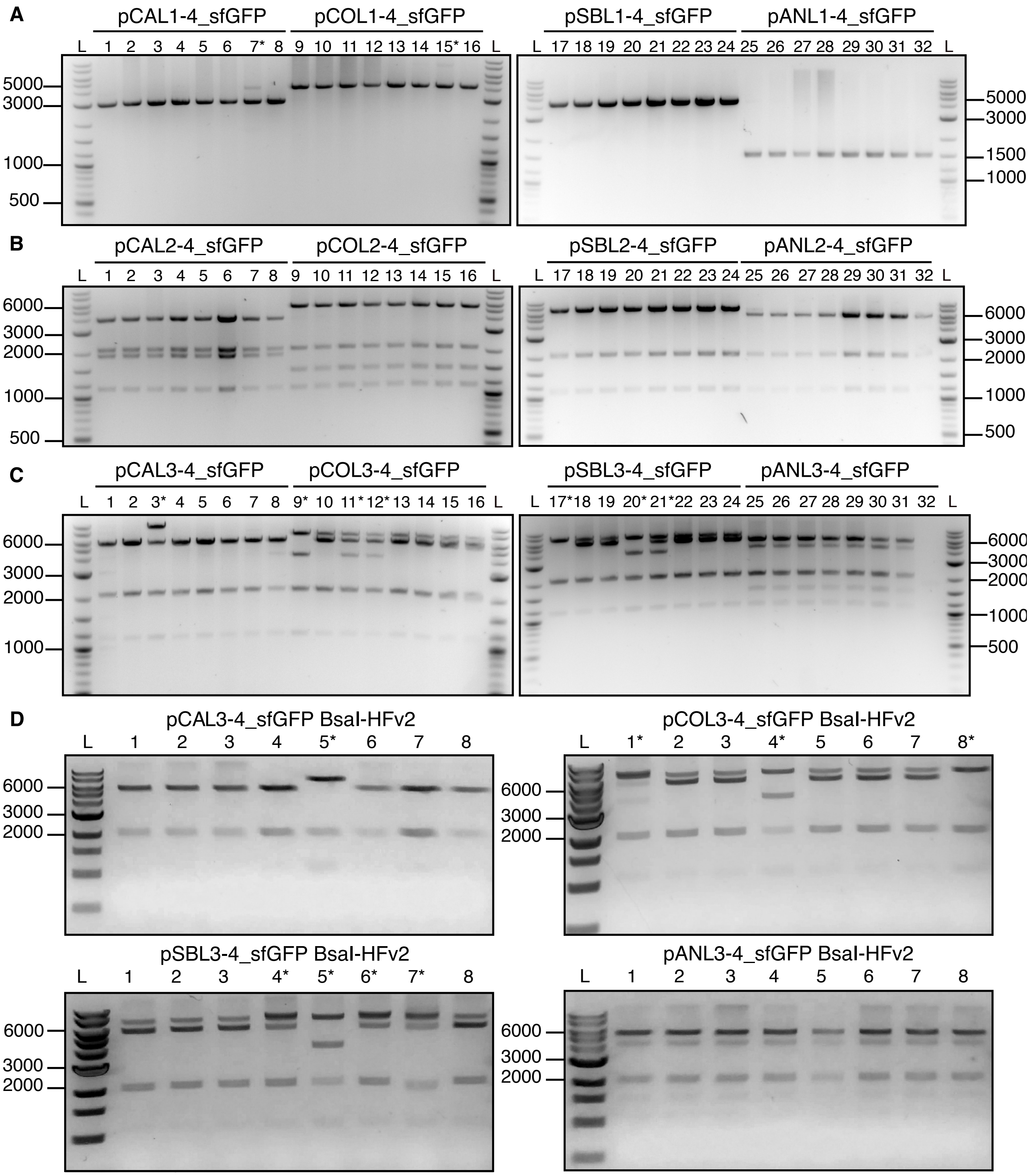

**Supplementary Figure S4. Integrity of assembly.**

Purified assemblies were evaluated by restriction profiling to determine integrity of assembly by comparison with expected digestion profiles. Incorrect assemblies are designated with an * next to the sample number. (A) Integrity of L1 assemblies. Purified L1 assemblies were digested with HindIII and restriction profiles determined by agarose-gel electrophoresis. (B) Integrity of L2 assemblies. Purified L2 assemblies were digested with XbaI and restriction profiles determined by agarose-gel electrophoresis. (C) Integrity of L3 assemblies. Purified L3 assemblies were digested with HindIII and restriction profiles determined by agarose-gel electrophoresis. (D) Integrity of L3 assemblies using BsaI-HFv2. Purified L3 assemblies generated with BsaI-HFv2 were digested with HindIII and restriction profiles determined by agarose-gel electrophoresis.

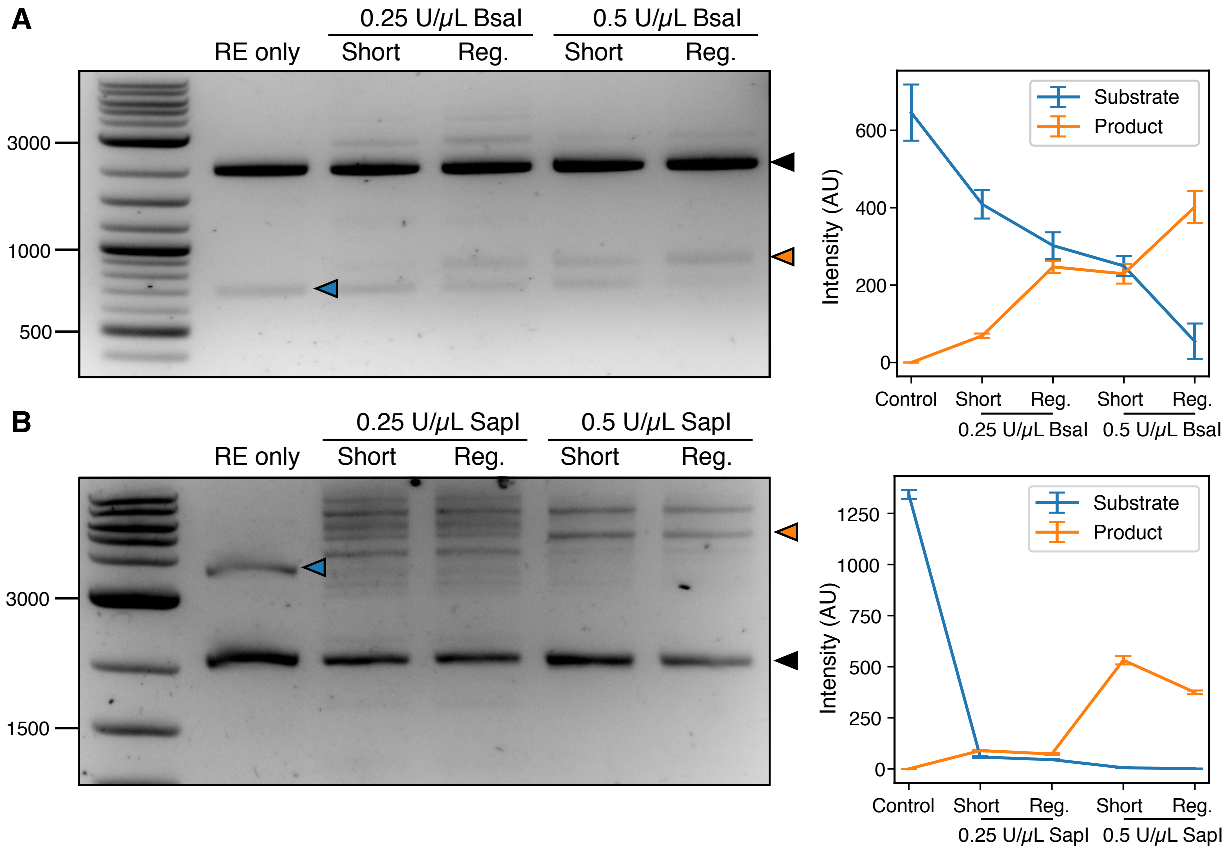

**Supplementary Figure S5. Reaction setup tests.**
(A) BsaI tests. An image for a resulting agarose gel electrophoresis of BsaI concentration and cycling condition tests is shown along with a lineplot with the quantification of the expected substrate and product bands of the reaction. The black arrow indicates the cut vector backbone, the blue arrow indicates the substrate and the orange arrow indicates the expected full-length product. (B) SapI tests. An image for a resulting agarose gel electrophoresis of BsaI concentration and cycling condition tests is shown along with a lineplot with the quantification of the expected substrate and product bands of the reaction. The black arrow indicates the cut vector backbone, the blue arrow indicates the substrate and the orange arrow indicates the expected full-length product.

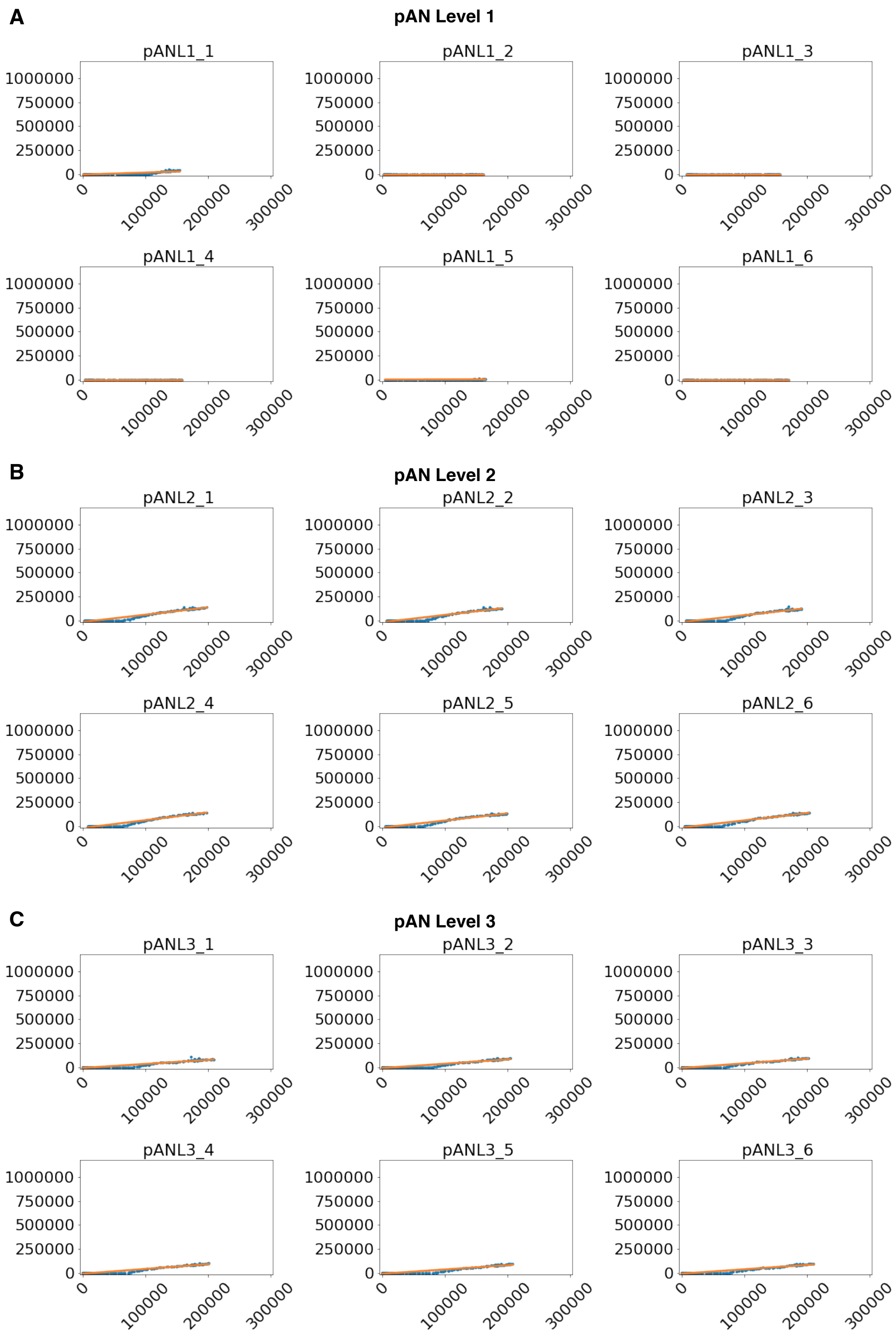

**Supplementary Figure S6. Representative replicates for performance evaluation of pAN plasmids in *E. coli*.** Plots showing GFP (y axis) against RFP (x axis) signal in arbitrary units of L1 (A), L2 (B) and L3 (C) assemblies. Green and red fluorescent signal correspond to plate fluorometry assays of the expression of a plasmid-borne GFP cassette and a chromosomally-integrated mRFP1 cassette.

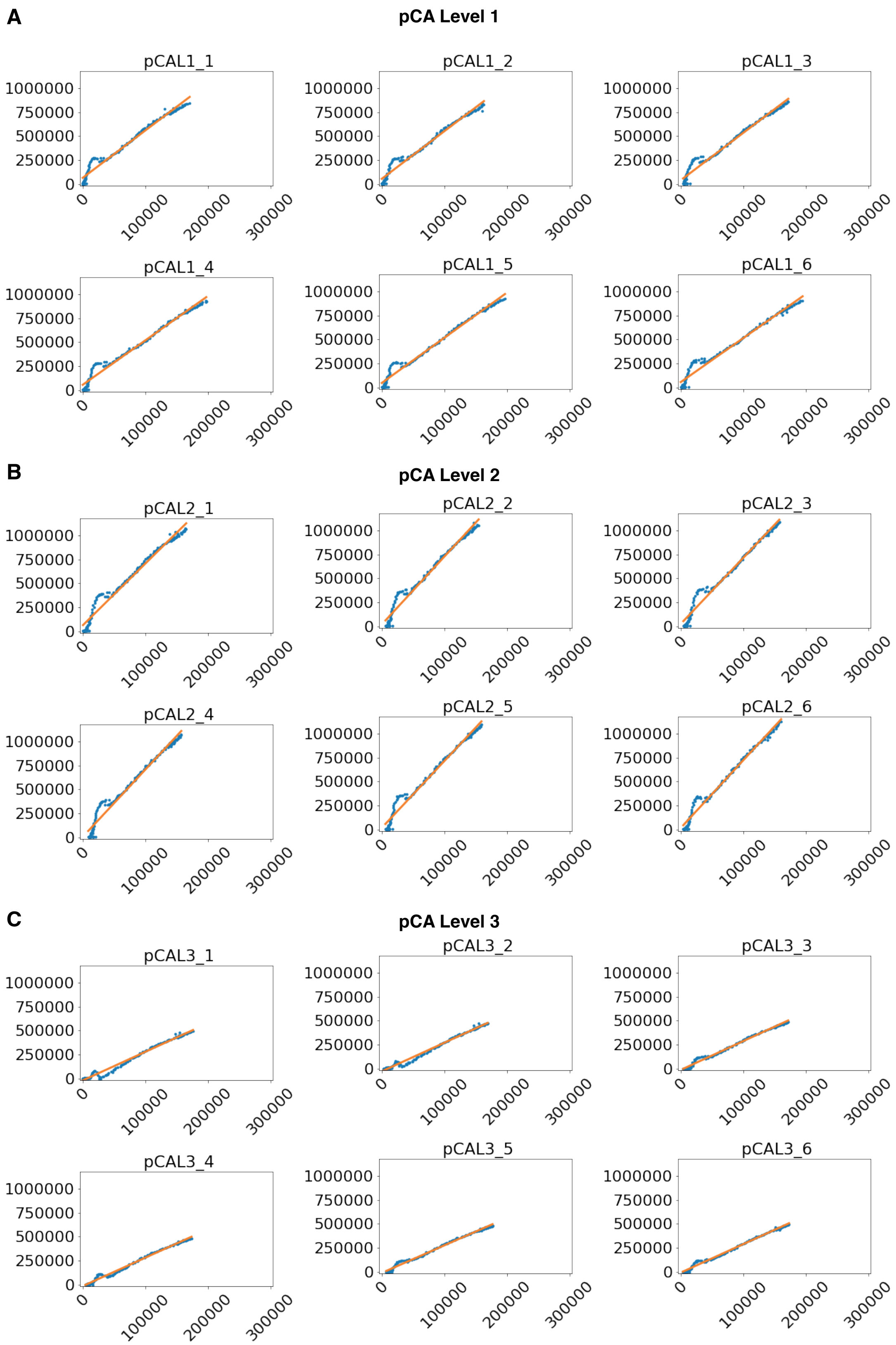

**Supplementary Figure S7. Representative replicates for performance evaluation of pCA plasmids in *E. coli*.** Plots showing GFP (y axis) against RFP (x axis) signal in arbitrary units of L1 (A), L2 (B) and L3 (C) assemblies. Green and red fluorescent signal correspond to plate fluorometry assays of the expression of a plasmid-borne GFP cassette and a chromosomally-integrated mRFP1 cassette.

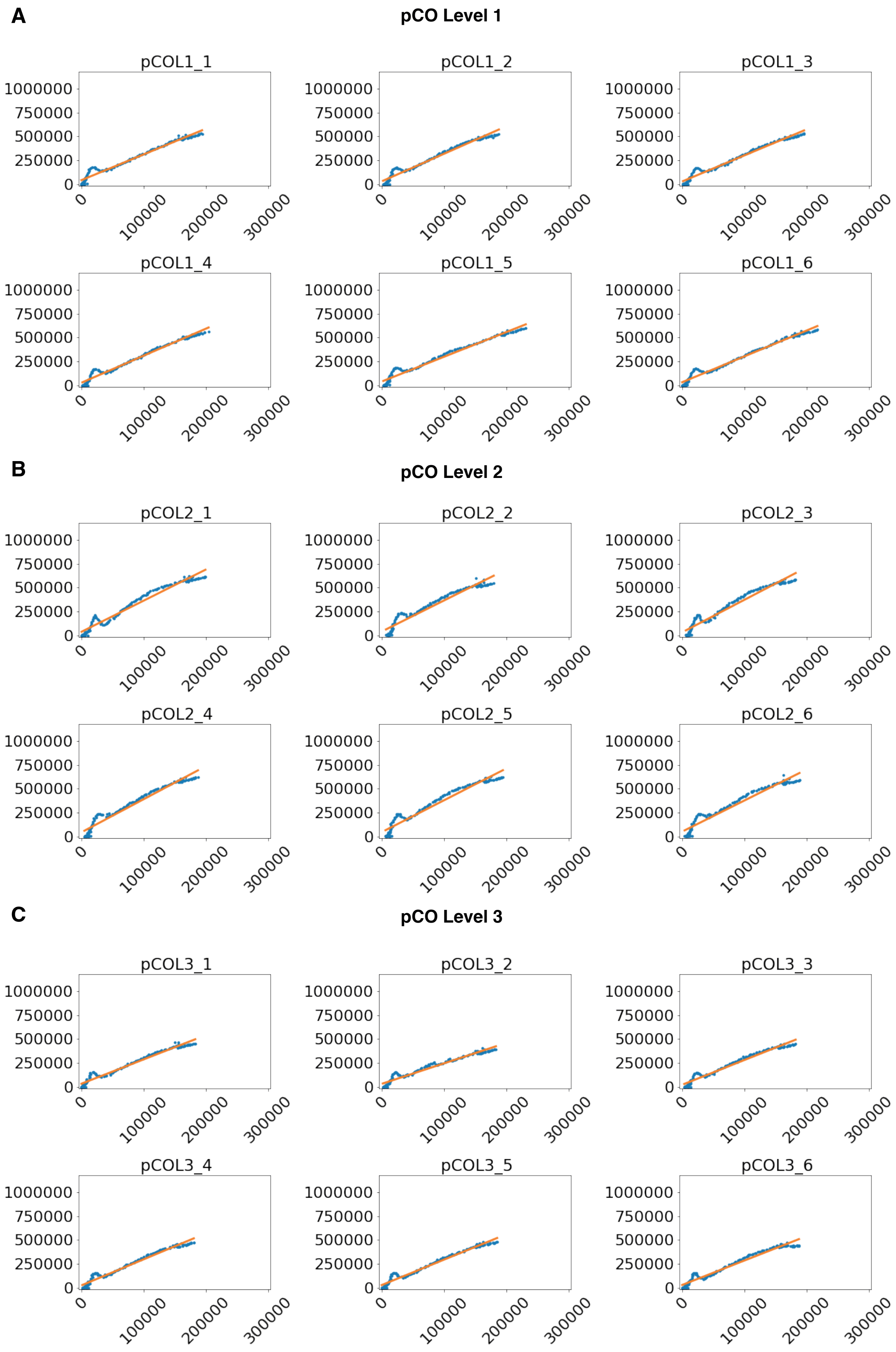

**Supplementary Figure S8. Representative replicates for performance evaluation of pCO plasmids in *E. coli*.** Plots showing GFP (y axis) against RFP (x axis) signal in arbitrary units of L1 (A), L2 (B) and L3 (C) assemblies. Green and red fluorescent signal correspond to plate fluorometry assays of the expression of a plasmid-borne GFP cassette and a chromosomally-integrated mRFP1 cassette.

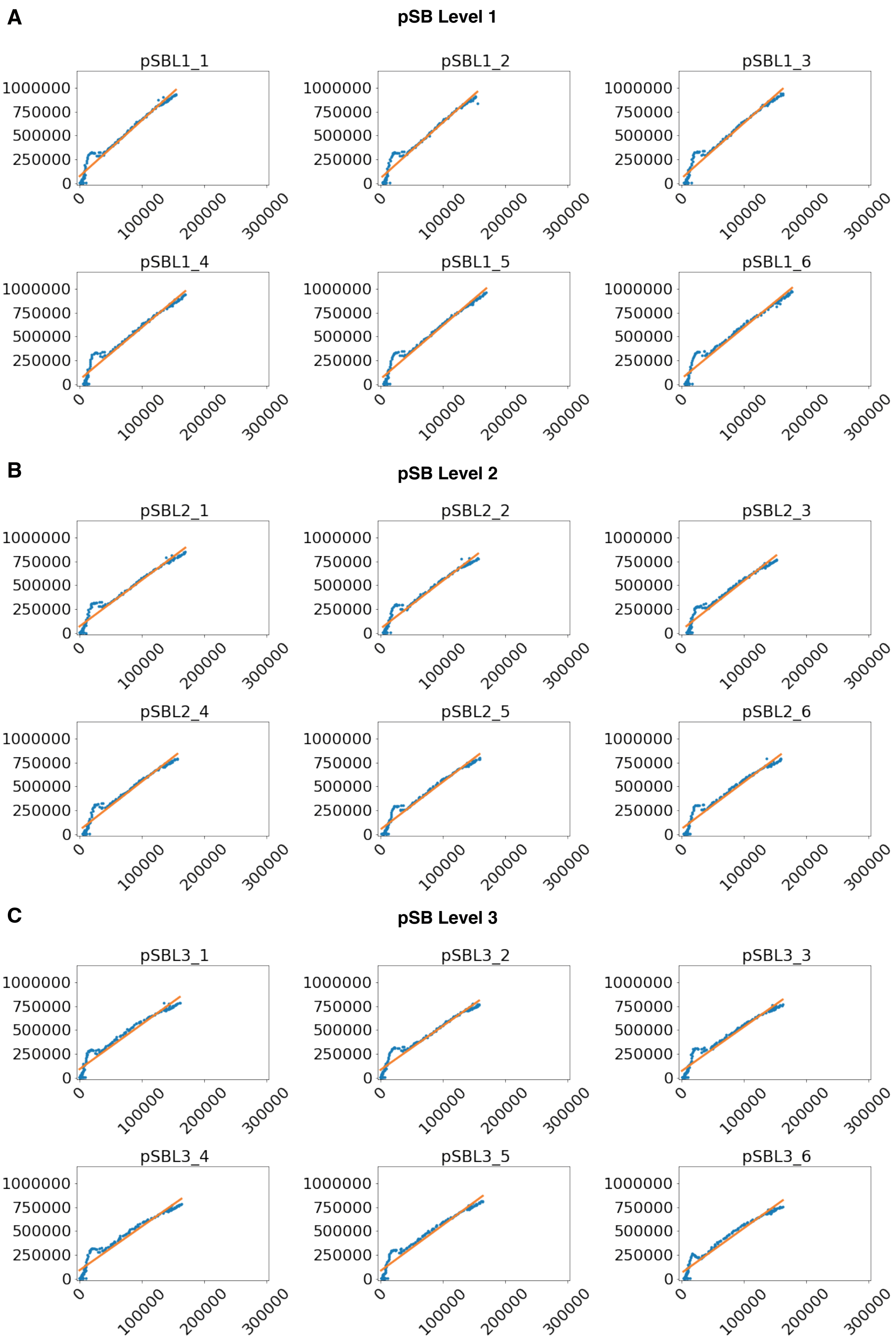

**Supplementary Figure S9. Representative replicates for performance evaluation of pSB plasmids in *E. coli*.** Plots showing GFP (y axis) against RFP (x axis) signal in arbitrary units of L1 (A), L2 (B) and L3 (C) assemblies. Green and red fluorescent signal correspond to plate fluorometry assays of the expression of a plasmid-borne GFP cassette and a chromosomally-integrated mRFP1 cassette.

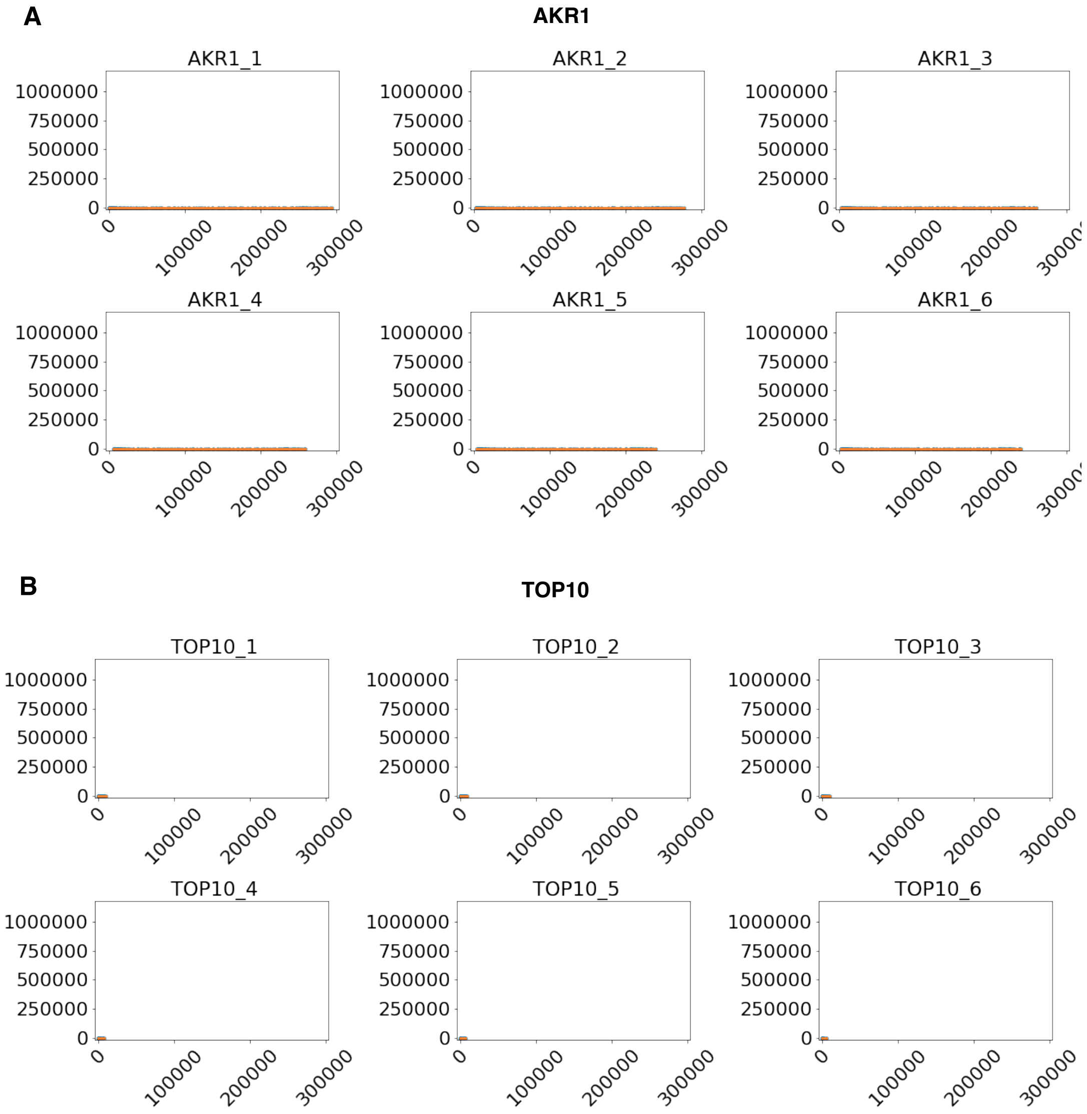

**Supplementary Figure S10. Representative replicates for performance evaluation controls.** AKR1(A) and TOP10 (B) controls *E. coli*. Fluorescence signal is in arbitrary units.

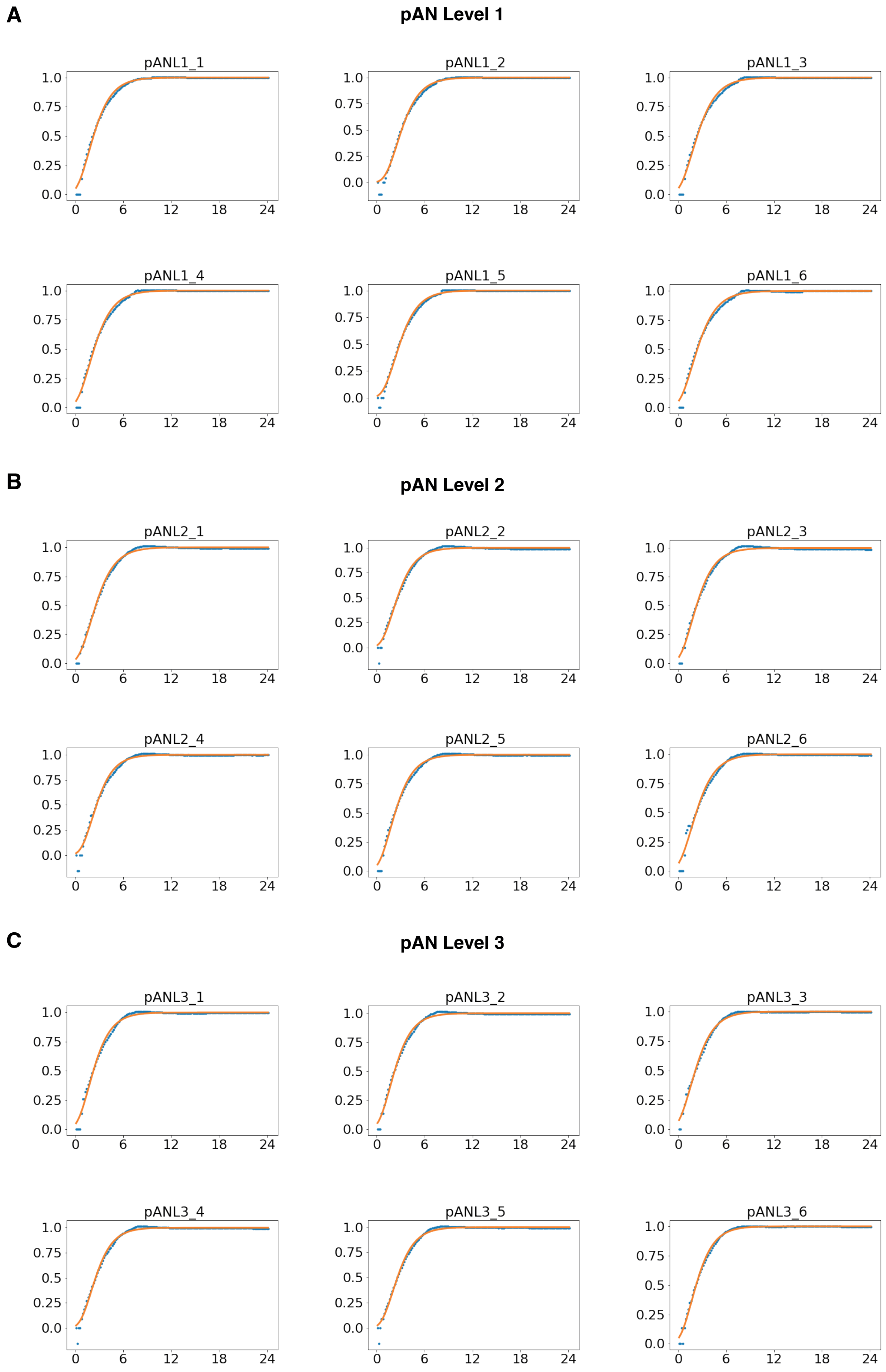

**Supplementary Figure S11. Representative replicates of growth effect evaluation of pAN plasmids in *E. coli*.** Plots showing normalised growth curves of L1 (A), L2 (B) and L3 (C) assemblies.

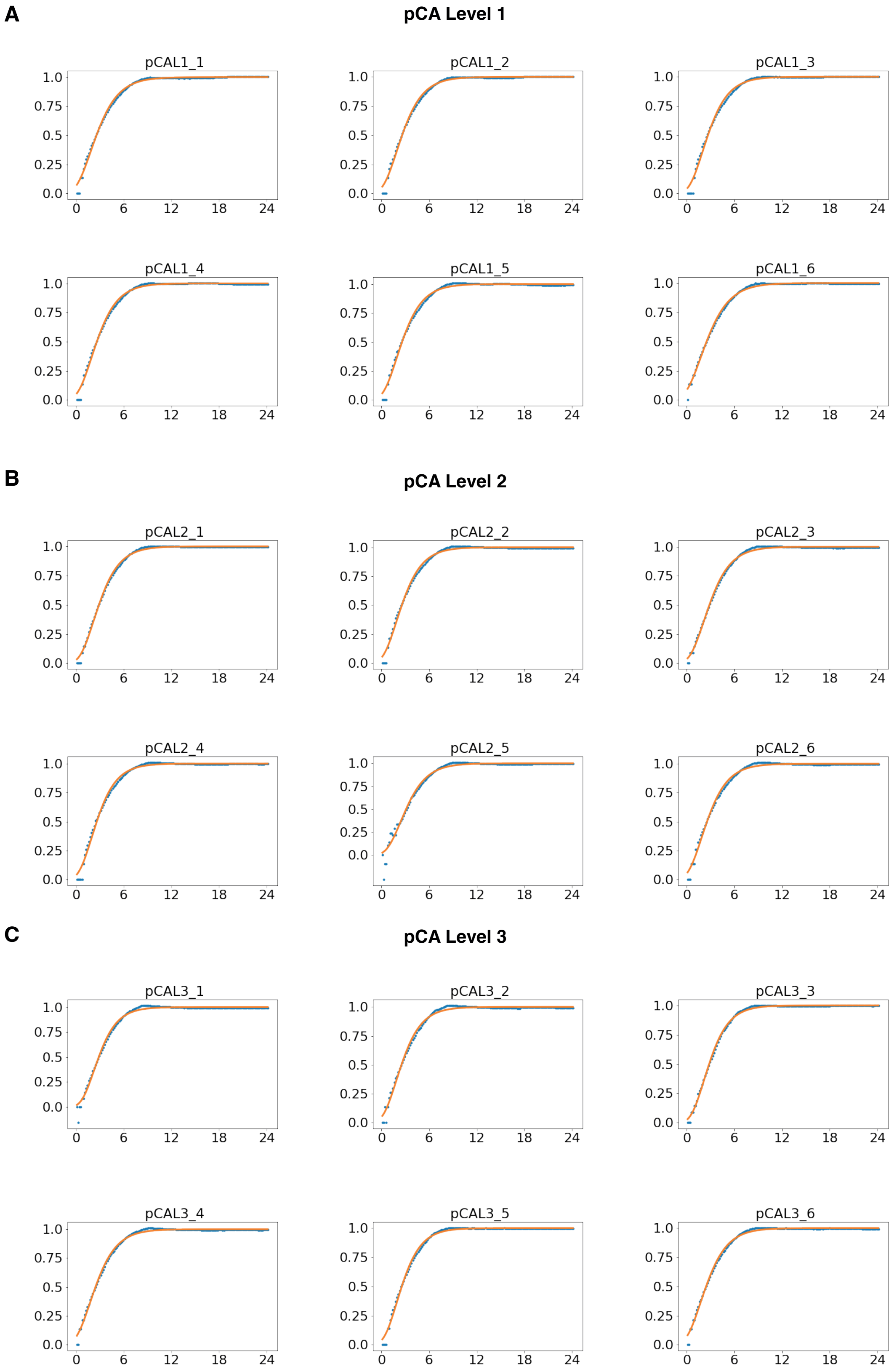

**Supplementary Figure S12. Representative replicates of growth effect evaluation of pCA plasmids in *E. coli*.** Plots showing normalised growth curves of L1 (A), L2 (B) and L3 (C) assemblies.

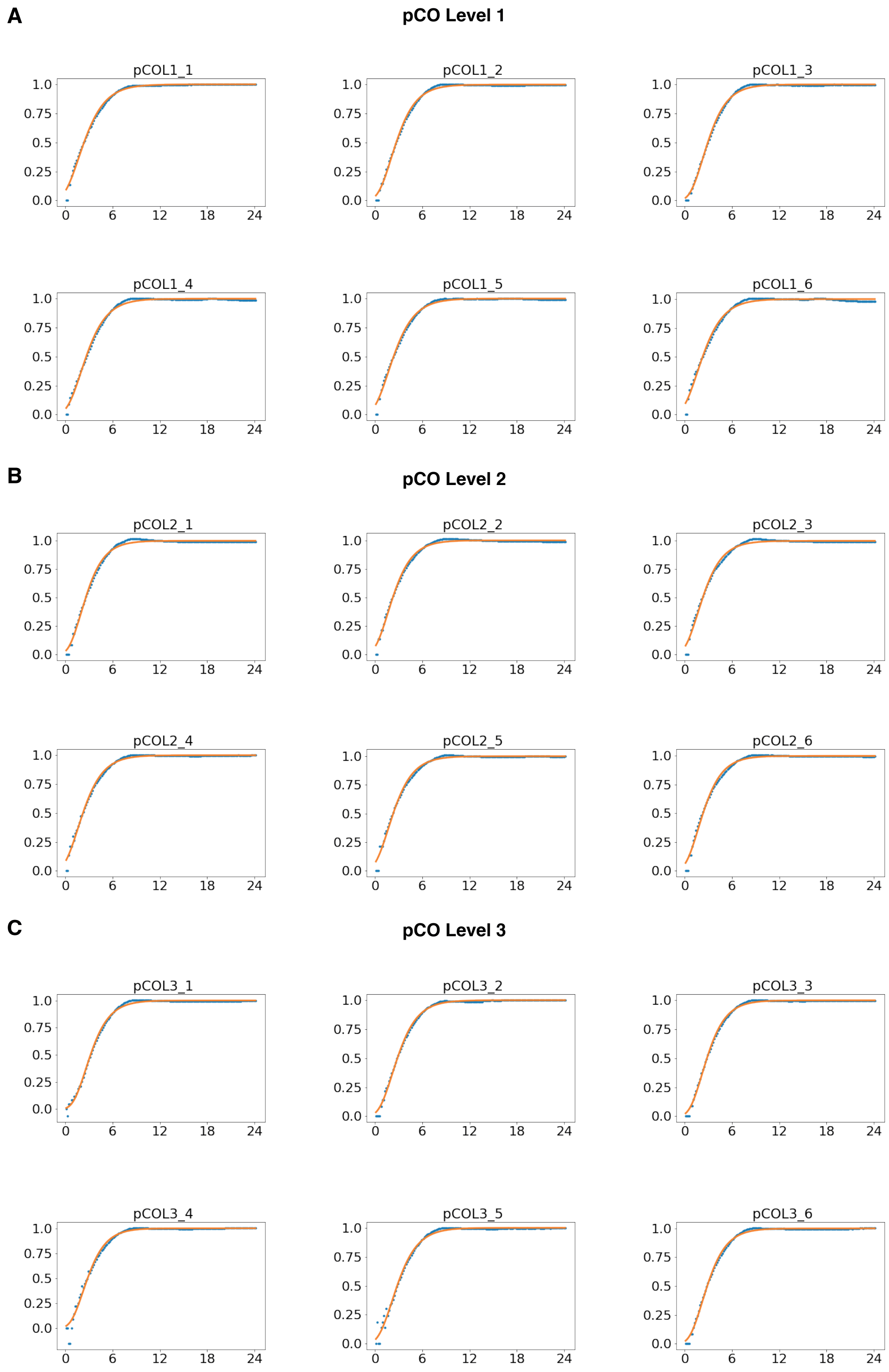

**Supplementary Figure S13. Representative replicates of growth effect evaluation of pCO plasmids in *E. coli*.** Plots showing normalised growth curves of L1 (A), L2 (B) and L3 (C) assemblies.

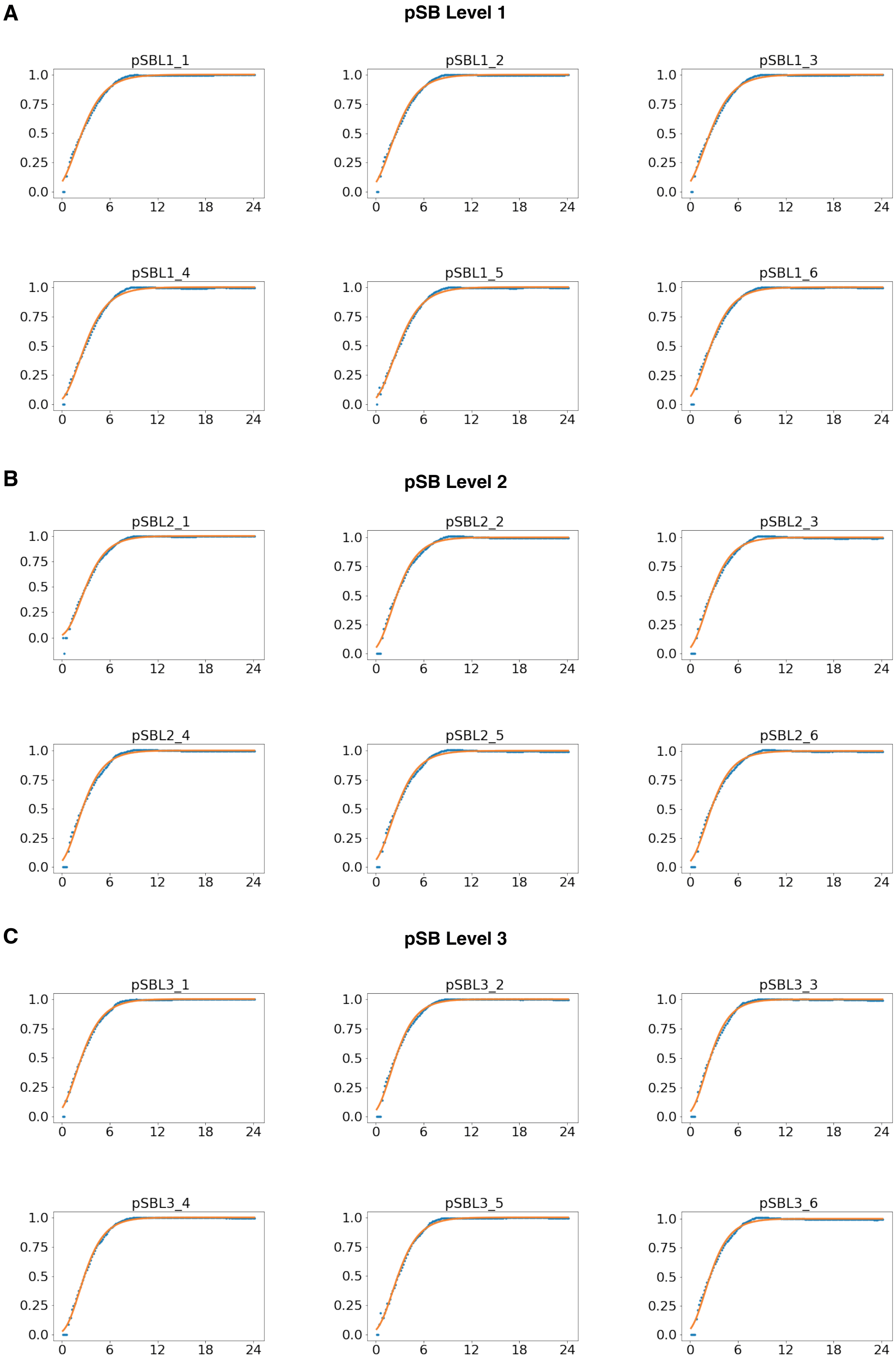

**Supplementary Figure S14. Representative replicates of growth effect evaluation of pSB plasmids in *E. coli*.** Plots showing normalised growth curves of L1 (A), L2 (B) and L3 (C) assemblies.

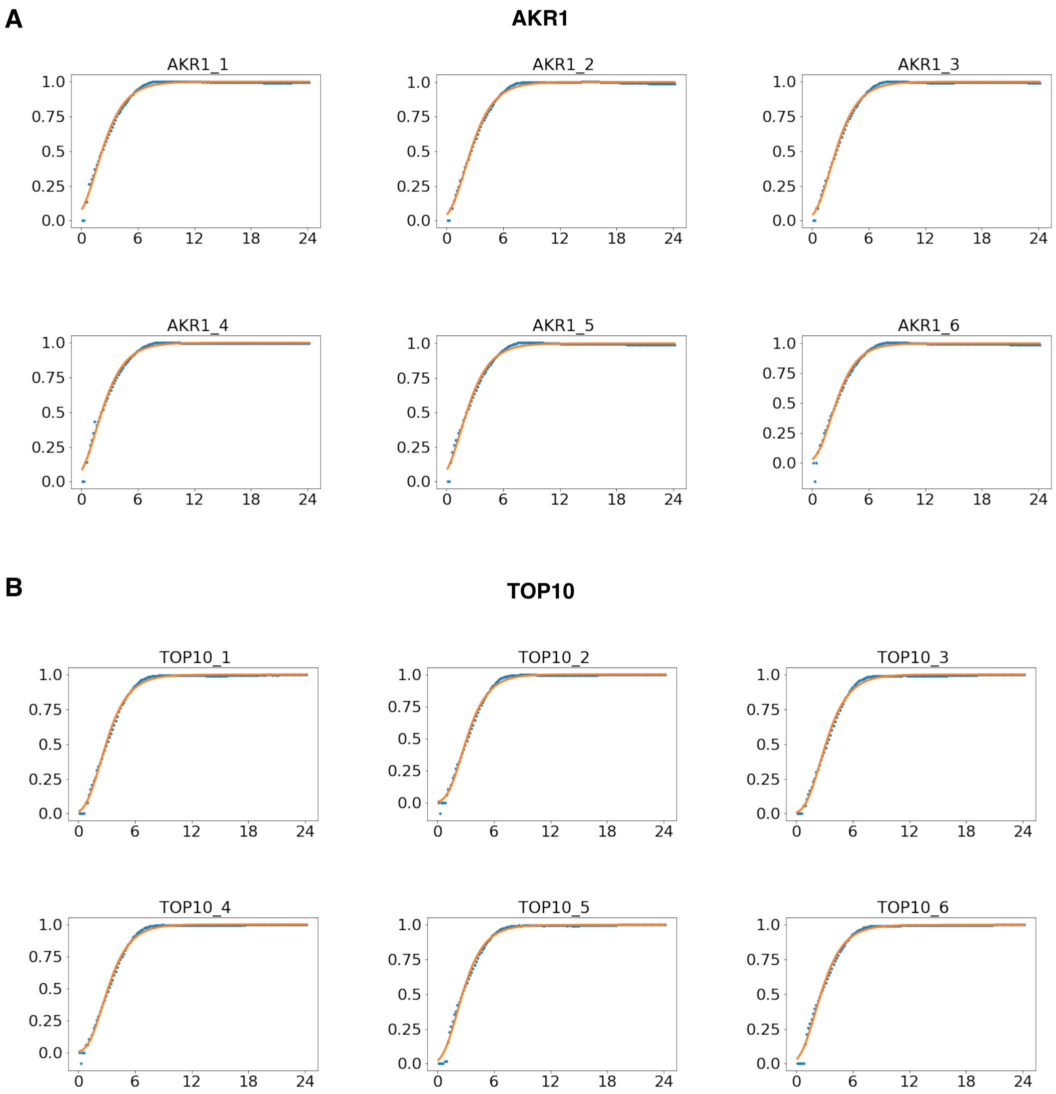

**Supplementary Figure S15. Representative replicates of growth effect evaluation controls.** *E.coli* control strains AKR1 (A) and TOP10 (B). Normalised growth rate values are shown.

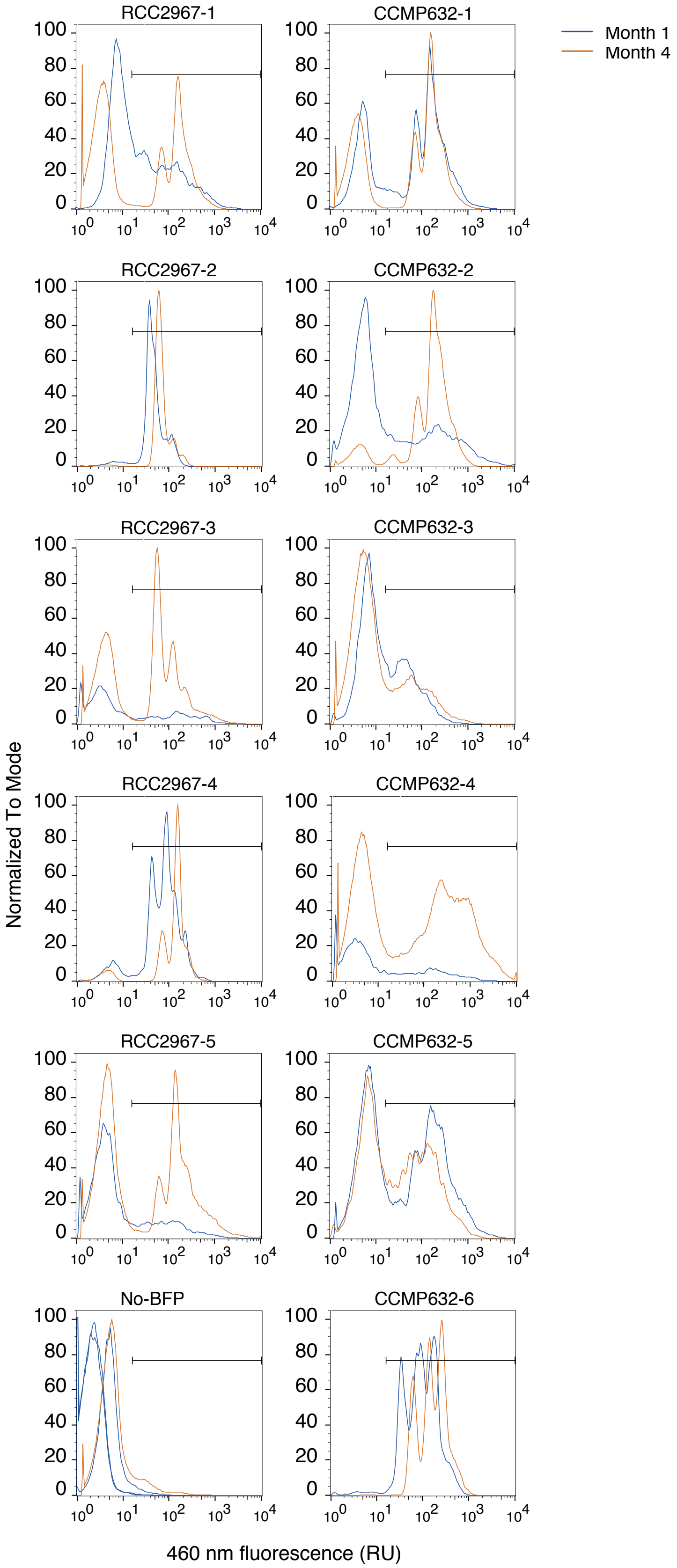

**Supplementary Figure S16. Stability of expression of uLoop plasmid in *P.tricornutum*.**
Histograms of blue fluorescence (exc. 405 nm, em. 460/30 nm) of ex-conjugates of *P. tricornutum* sub-strains RCC2967 and CCMP632 expressing mTagBFP2 under the control of the H4 promoter, tested after first growth in liquid medium (month 1) and three months later (month 4). Bar shows the gate defining blue fluorescent cells, with the lower limit based on the maximum 95th percentile of blue fluorescence in no-BFP controls (exconjugates with a plasmid containing the nonfluorescent luxR gene) run in parallel with the same cytometer settings.

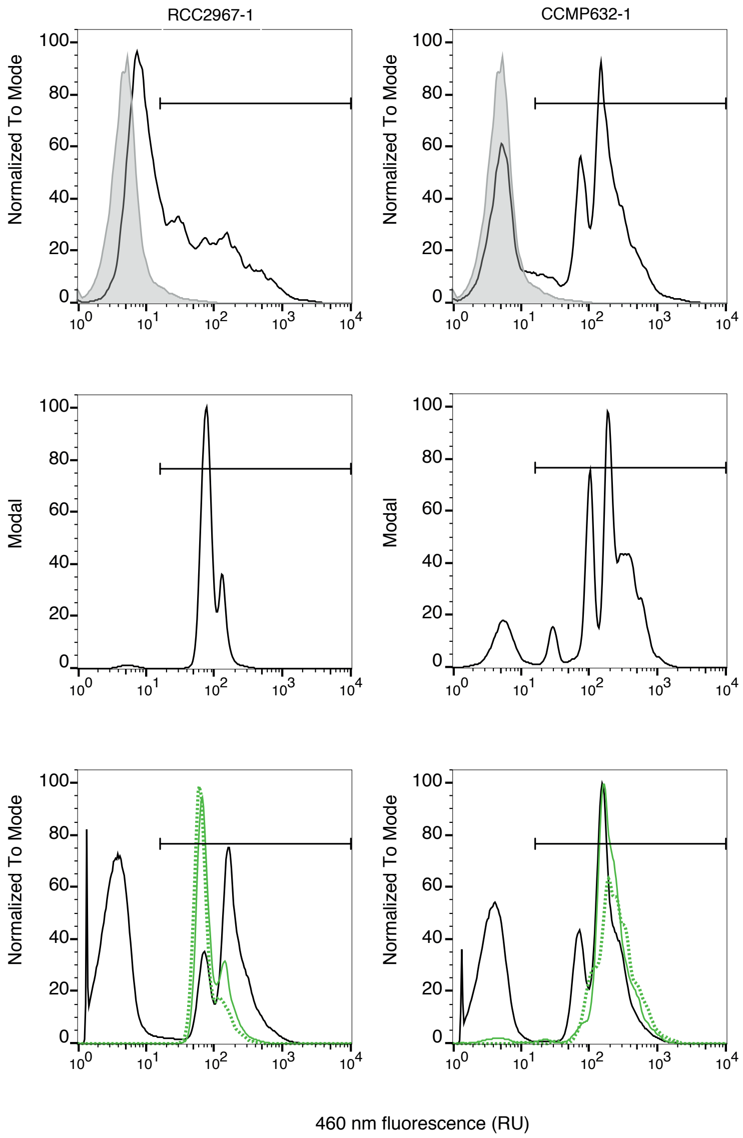

**Supplementary Figure S17. Histograms of** ***P.tricornutum*** **ex-conjugate fluorescence before and after cell sorting.** Top two panels: Histograms at month 0, without sorting. Also shown for comparison is the histogram of ex-conjugates with the gene for the non-fluorescent protein LuxR (light grey). Middle two panels: 2.5 months after conjugation, when blue-fluorescent cells were sorted. Bottom two panels: 4 months after conjugation, comparing cultures from sorted cells (green) to original ex-conjugates (black), both allowed to evolve through several transfers in liquid medium with zeocin. Bar shows the gate defining blue fluorescent cells, with the lower limit based on the maximum 95th percentile of blue fluorescence in No-BFP controls (exconjugates with a plasmid containing the nonfluorescent luxR gene) run in parallel with the same cytometer settings.

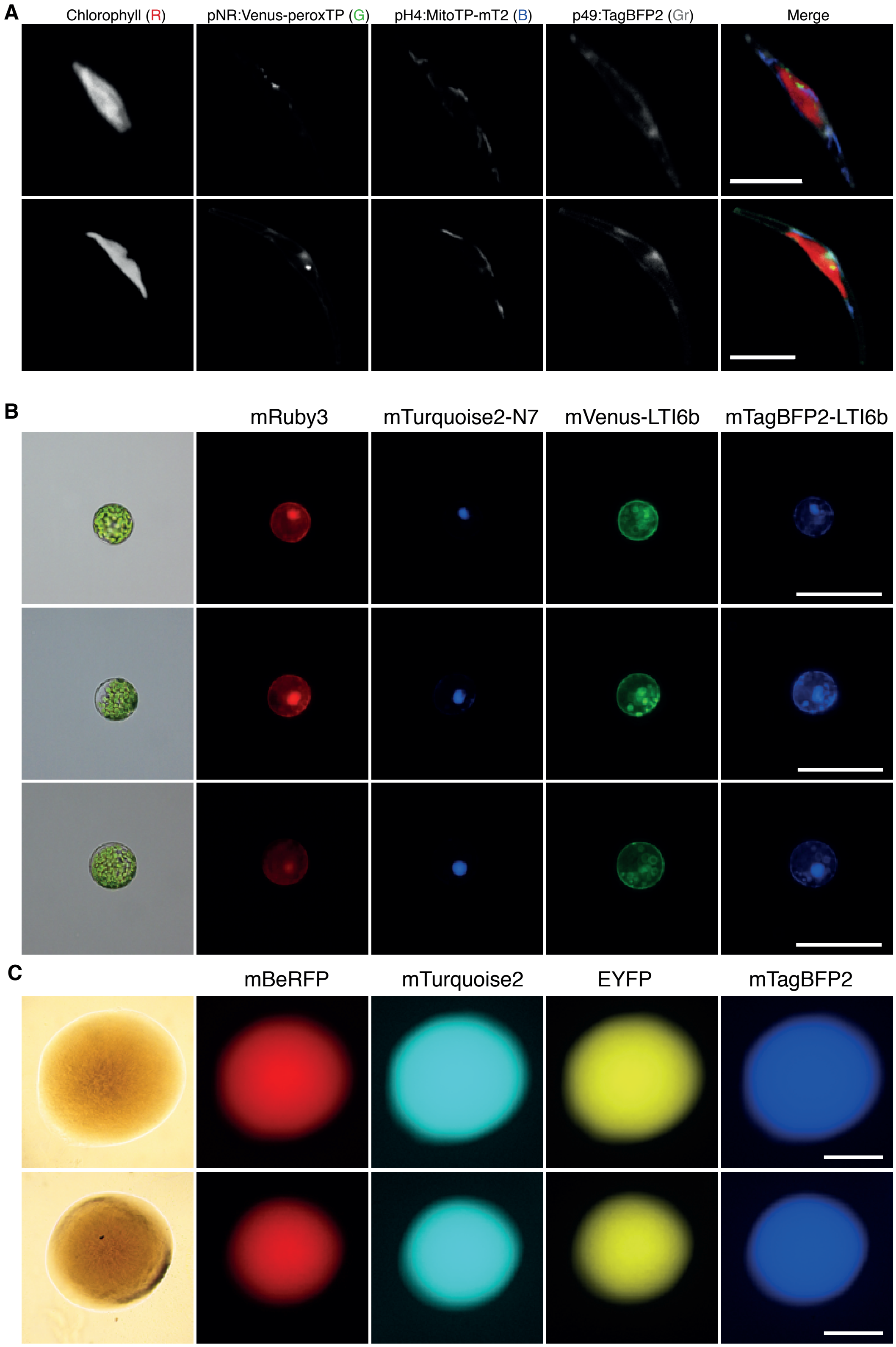

**Supplementary Figure S18. Use of uLoop vectors across multiple organisms.**
(A) Two representative individuals of *P. tricornutum* exconjugants transformed with pCAL2-1-FPrep showing expression of three fluorescent reporters. From left to right, Chlorophyll fluorescence, mVenus fluorescent protein fused to a peroxisomal localisation tag, mTurquoise2 fluorescent protein fused to a mitochondrial localisation tag and mTagBFP2 fluorescent protein expressed in the cytoplasm. Scale bar = 10 $\mu$m. (B) Three representative protoplasts of *Arabidopsis thaliana* transformed with pSBL2-1-4xFP vector harboring four fluorescent reporters. From left to right, mRuby3 fluorescent protein expressed in the cytoplasm, mTurquoise2 fluorescent protein fused to the nuclear localisation tab N7, mVenus fluorescent protein fused to plasma-membrane localisation signal LTi6b and mTagBFP2 fluorescent protein fused to plasma-membrane localisation signal LTi6b. Scale bar = 100 $\mu$m. (C) Expression of four fluorescent reporters in *Escherichia coli* colonies transformed with showing expression of mBeRFP, mTurquoise2, EYFP and mTagBFP2 fluorescent proteins pCA (top) and pSB (bottom) uLoop vectors. Scale bar = 500 $\mu$m.

| Plasmid | Level | Mean | STD | SEM | 95% CI |
|---|---|---|---|---|---|
| pCA | Level 1 | 4,43 | 0,53 | 0,31 | [3,12 - 5,75] |
| pCA | Level 2 | 5,81 | 1,01 | 0,58 | [3,29 - 8,32] |
| pCA | Level 3 | 2,57 | 0,42 | 0,25 | [1,51 - 3,62] |
| pCO | Level 1 | 2,28 | 0,50 | 0,29 | [1,04 - 3,52] |
| pCO | Level 2 | 2,80 | 0,54 | 0,31 | [1,46 - 4,13] |
| pCO | Level 3 | 2,05 | 0,44 | 0,25 | [0,97 - 3,13] |
| pSB | Level 1 | 5,02 | 0,75 | 0,43 | [3,15 - 6,88] |
| pSB | Level 2 | 4,33 | 0,65 | 0,37 | [2,72 - 5,94] |
| pSB | Level 3 | 4,18 | 0,49 | 0,28 | [2,97 - 5,39] |
| pAN | Level 1 | 0,03 | 0,03 | 0,02 | [-0,04 - 0,10] |
| pAN | Level 2 | 0,41 | 0,31 | 0,18 | [-0,37 - 1,18] |
| pAN | Level 3 | 0,23 | 0,20 | 0,12 | [-0,27 - 0,73] |
| AKR1 | - | 0,00 | 0,00 | 0,00 | [0,00 - 0,00] |

**Supplementary Table S1. Green-to-red fluorescent ratio values for *E.coli* plate fluorometry.**
The mean, standard deviation (STD), standard error of the mean (SEM) and 95% confidence intervals are shown for each plasmid level and group. AKR1 does not contain 'Level' plasmids, thus only the group value is reported. Values correspond to three independent replicates (N=3).

| | DF | SS | MSS | F | PR(>F) |
|---|---|---|---|---|---|
| **Level** | 2.0 | 7.141522 | 3.570761 | 0.941299 | 0.400345 |
| **Residual** | 33.0 | 125.183476 | 3.793439 | | |

**Supplementary Table S2. ANOVA test for *E.coli* plate fluorometry Green/Red fluorescent ratio values between levels across the four kits (pCA, pSB, pCO, pAN).**
A two-way ANOVA comparison was performed for L1, L2 and L3 values of the four kits (pCA, pSB, pCO, pAN) grouped together per level. There were no statistically significant differences between group means (i.e. plasmid sizes). DF: degrees of freedom; SS: sum of squares; MSS: mean sum of squares; F: F statistic value; PR(>F): p-value of F statistic. AKR1 data was not considered.

| Plasmid set | | DF | SS | MSS | F | PR(>F) |
|---|---|---|---|---|---|---|
| **pCA** | **Level** | 2.0 | 15.866.420 | 793.321 | 16.008.907 | 0.003931 |
| | **Residual** | 6.0 | 2.973.299 | 0.49555 | | |
| **pCO** | **Level** | 2.0 | 0.876727 | 0.438363 | 1.803.301 | 0.243638 |
| | **Residual** | 6.0 | 1.458.537 | 0.243089 | | |
| **pSB** | **Level** | 2.0 | 1.187.523 | 0.593761 | 1.461.676 | 0.303997 |
| | **Residual** | 6.0 | 2.437.317 | 0.406220 | | |
| **pAN** | **Level** | 2.0 | 0.213126 | 0.106563 | 2.301.536 | 0.1812 |
| | **Residual** | 6.0 | 0.277804 | 0.046301 | | |

**Supplementary Table S3. ANOVA test for E. coli plate fluorometry Green/Red fluorescent ratio values comparing levels within each plasmid kit.**
A two-way ANOVA comparison was performed among L1, L2 and L3 values for each plasmid kit. There were statistically significant differences between levels of pCA kit. DF: degrees of freedom; SS: sum of squares; MSS: mean sum of squares; F: F statistic value; PR(>F): p-value of F statistics.

| Group 1 | Group 2 | Mean Diff | Lower | Upper | p-value | Reject |
|---|---|---|---|---|---|---|
| Level 1 | Level 2 | 1.37 | -0.39 | 3.13 | 0.1174 | False |
| Level 1 | Level 3 | -1.87 | -3.63 | -0.11 | 0.0399 | True |
| Level 2 | Level 3 | -3.24 | -5.00 | -1.48 | 0.0032 | True |

**Supplementary Table S4. pCA multiple comparison of means (Tukey HSD test) for E. coli plate fluorometry Green/Red fluorescent ratio values.**
Using rejection under 95% significance for each pair compared, showed statistically significant differences for level 3 of pCA.

| Plasmid set | | DF | SS | MSS | F | PR(>F) |
|---|---|---|---|---|---|---|
| pCA | Level | 2 | 2.401 | 1.200 | 11.491 | 0.009 |
| | Residual | 6 | 0.627 | 0.104 | | |
| pCO | Level | 2 | 0.149 | 0.074 | 1.115 | 0.387 |
| | Residual | 6 | 0.400 | 0.067 | | |
| pSB | Level | 2 | 0.4756 | 0.2378 | 2.6964 | 0.1461 |
| | Residual | 6 | 0.5292 | 0.0882 | | |
| pAN | Level | 2 | 0.3474 | 0.1737 | 12.6725 | 0.0070 |
| | Residual | 6 | 0.0822 | 0.0137 | | |

**Supplementary Table S5. Internal group ANOVA test for *E. coli* flow cytometry Green/Red fluorescent ratio values.**
A two-way ANOVA comparison was performed among L1, L2 and L3 values for each plasmid kit. There were statistically significant differences between uLoop levels of pCA and pAN vector kits. DF: degrees of freedom; SS: sum of squares; MSS: mean sum of squares; F: F statistic value; PR(>F): p-value of F statistic.

| Vector Set | Group 1 | Group 2 | Mean Diff | Lower | Upper | p-value | Reject |
|---|---|---|---|---|---|---|---|
| **pAN** | pANL1 | pANL2 | 0,401 | 0,108 | 0,694 | 0,0135 | **True** |
| | pANL1 | pANL3 | -0,030 | -0,323 | 0,263 | 0,9000 | False |
| | pANL2 | pANL3 | -0,431 | -0,724 | -0,138 | 0,0096 | **True** |
| **pCA** | pCAL1 | pCAL2 | 0,239 | -0,570 | 1,047 | 0,6521 | False |
| | pCAL1 | pCAL3 | -0,957 | -1,766 | -0,148 | 0,0255 | **True** |
| | pCAL2 | pCAL3 | -1,195 | -2,004 | -0,387 | 0,0094 | **True** |

**Supplementary Table S6. pAN and pCA multiple comparison of means (Tukey HSD test) for *E.coli* flow cytometry Green/Red fluorescent ratio values.**
Rejection was performed under 95% significance for each pair compared.

| | | Plate fluorometry | | | Flow cytometry | | |
|---|---|---|---|---|---|---|---|
| **Plasmid** | **Level** | **Mean** | **STD** | **CV** | **Mean** | **STD** | **CV** |
| pCA | All | 4,27 | 1,53 | 35,95 | 1,50 | 0,62 | 40,99 |
| pCO | All | 2,37 | 0,54 | 22,75 | 1,37 | 0,26 | 19,16 |
| pSB | All | 4,51 | 0,67 | 14,93 | 1,67 | 0,35 | 21,28 |
| pAN | All | 0,22 | 0,25 | 111,31 | 0,57 | 0,23 | 40,55 |

**Supplementary Table S7. Green-to-red fluorescent ratio values for *E.coli* plate fluorometry and flow cytometry by plasmid group.**
The mean, standard deviation (STD), and coefficient of variation (CV) are shown for each vector group. Values correspond to three independent replicates and three levels per group (N=9).

| Plasmid | Level | Mean | STD | SEM | 95% CI |
|---|---|---|---|---|---|
| pCA | Level 1 | 1,26 | 0,20 | 0,12 | [0,76 - 1,76] |
| pCA | Level 2 | 1,25 | 0,14 | 0,08 | [0,89 - 1,60] |
| pCA | Level 3 | 1,33 | 0,15 | 0,09 | [0,94 - 1,71] |
| pCA | All | 1,28 | 0,15 | 0,05 | [1,16 - 1,39] |
| pCO | Level 1 | 1,35 | 0,05 | 0,03 | [1,23 - 1,46] |
| pCO | Level 2 | 1,34 | 0,19 | 0,11 | [0,86 - 1,82] |
| pCO | Level 3 | 1,42 | 0,17 | 0,10 | [1,01 - 1,83] |
| pCO | All | 1,37 | 0,14 | 0,05 | [1,26 - 1,47] |
| pSB | Level 1 | 1,17 | 0,14 | 0,08 | [0,81 - 1,52] |
| pSB | Level 2 | 1,32 | 0,20 | 0,12 | [0,81 - 1,83] |
| pSB | Level 3 | 1,25 | 0,22 | 0,13 | [0,70 - 1,79] |
| pSB | All | 1,24 | 0,18 | 0,06 | [1,11 - 1,38] |
| pAN | Level 1 | 1,43 | 0,12 | 0,07 | [1,13 - 1,73] |
| pAN | Level 2 | 1,37 | 0,20 | 0,12 | [0,87 - 1,87] |
| pAN | Level 3 | 1,47 | 0,22 | 0,13 | [0,93 - 2,01] |
| pAN | All | 1,42 | 0,17 | 0,06 | [1,30 - 1,55] |
| AKR1 | - | 1,53 | 0,18 | 0,10 | [0,18 - 0,10] |

**Supplementary Table S8. *E. coli* growth rate values.**
Mean, standard deviation (STD), standard error of the mean(SEM) and 95% confidence intervals are shown for each plasmid level and group. AKR1 does not contain 'Level' plasmids, thus only the group value is reported. Values correspond to three whole independent replicates (N=3).

| Plasmid set | | DF | SS | MSS | F | PR(>F) |
|---|---|---|---|---|---|---|
| pCA | Level | 2.0 | 0.011597 | 0.005798 | 0.203773 | 0.821067 |
| | Residual | 6.0 | 0.170733 | 0.028456 | | |
| pCO | Level | 2.0 | 0.012275 | 0.006138 | 0.272106 | 0.770694 |
| | Residual | 6.0 | 0.135338 | 0.022556 | | |
| pSB | Level | 2.0 | 0.035281 | 0.01764 | 0.480012 | 0.640651 |
| | Residual | 6.0 | 0.220500 | 0.03675 | | |
| pAN | Level | 2.0 | 0.013763 | 0.006881 | 0.201582 | 0.822754 |
| | Residual | 6.0 | 0.204824 | 0.034137 | | |

**Supplementary Table S9. ANOVA test for E. coli plate fluorometry growth rate values.**
A two-way ANOVA comparison was performed among L1, L2 and L3 values for each plasmid kit. There was no statistically significant differences. DF: degrees of freedom; SS: sum of squares; MSS: mean sum of squares; F: F statistic value; PR(>F): p-value of F statistic.

| N | Level | Pos. | Name | Description | Notes | Submitter/Source |
|---|---|---|---|---|---|---|
| 1 | L0 | AB | AB_J23101 | J23101 promoter (E.coli) | L0>L1 Rx control | Matute & Núñez/[4] |
| 2 | L0 | AB | AB_R0010 (pLac) | R0010 promoter (MoClo: C1) | AmpR | [1] |
| 3 | L0 | AC | AC_CEN_ARS_HIS | Yeast centromere (PtPBR) | | Vince Bielinski |
| 4 | L0 | AC | AC_p49202 | Pt-p49202 | | Patrick Brunson |
| 5 | L0 | AC | AC_pFcpB | Pt-pFcpB | | Patrick Brunson |
| 6 | L0 | AC | AC_pH4 | Pt-pH4 | | Patrick Brunson |
| 7 | L0 | AC | AC_pNR | Pt-pNR | | Patrick Brunson |
| 8 | L0 | AC | AC_pUBQ10 | ubiqutin-10 gene promoter (A. thaliana) | | Ariel Cerda [5] |
| 9 | L0 | AC | AC_pTDH3 | pTDH3 promoter | SpectR | Matute & Núñez/[6] |
| 10 | L0 | AC | AC_pTEF1 | pTEF1 promoter | SpectR | Matute & Núñez/[6] |
| 11 | L0 | AF | AF_Spacer1 | L0 universal spacer 1 | | Bernardo Pollak |
| 12 | L0 | AF | AF_Spacer2 | L0 universal spacer 2 | | Bernardo Pollak |
| 13 | L0 | AF | AF_2u | 2u ori (yeast) | SpectR; Domestication from p426 (Addgene #43803; DiCarlo et al., 2013). | Matute & Núñez/[7] |
| 14 | L0 | AF | AF_CEN | CEN ori (yeast) | SpectR | Matute & Núñez/[6] |
| 15 | L0 | AF | AF_URA3 | Encodes URA3 | SpectR | Matute & Núñez/[6] |
| 16 | L0 | BC | BC-B0034m | BBa_B0034m RBS (E. coli) | L0>L1 Rx control | [1] |
| 17 | L0 | BC | BC_RiboJ54 | RiboJ54 F1RBS | AmpR | Matute & Núñez/[4] |
| 18 | L0 | CD | CD_mRFP1 | mRFP1 FP | | Bernardo Pollak/[8] |
| 19 | L0 | CD | CD_Venus | Venus FP | | Bernardo Pollak |
| 20 | L0 | CD | CD_MitoTP | Pt Mitochondrial targeting peptide | | Chris Dupont |
| 21 | L0 | CD | CD_mTagBFP2 | mTagBFP2, blue FP | STOP, AmpR. Chemical synthesis from sequence in Subach et al (2011). | Matute & Núñez/[9] |
| 22 | L0 | CD | CD_mTurquoise2 (yeast) | CDS mTurquoise2 (yeast) | SpectR | Matute & Núñez/[6] |
| 23 | L0 | CD | CD_ E0030 (YFP) | E0030 (MoClo: E4) | AmpR | [1] |
| 24 | L0 | CD | CD_mBeRFP | mBeRFP FP | AmpR | Matute & Núñez/[4] |
| 25 | L0 | CD | CD_sfGFP | sfGFP FP | SpectR | Bernardo Pollak |
| 26 | L0 | CD | CD_Tur2 | mTurquoise2 FP | AmpR | Matute & Núñez/[4] |
| 27 | L0 | CD | CD_LuxR | C0062 (MoClo: E1) | AmpR | [1] |
| 28 | L0 | CE | CE_OriTv2 | OriT v.2 | | Vince Bielinski |
| 29 | L0 | CE | CE_sfGFP | sfGFP FP, green FP | L0>L1 Rx control, STOP | Bernardo Pollak/[10] |
| 30 | L0 | CE | CE_mRuby3 | mRuby3 red FP | | Bernardo Pollak/[11] |
| 31 | L0 | DE | DE_mNeonGreen | mNeonGreen FP | | Bernardo Pollak |
| 32 | L0 | DE | DE_mTurquoise2 | mTurquoise2 cyan FP | | Bernardo Pollak/ [10] |
| 33 | L0 | DE | DE_sfGFP | sfGFP FP, green FP | | Bernardo Pollak/[10] |
| 34 | L0 | DE | DE_Venus-ThrHISFLAG | Venus YFP with ThrHisFLAG | | Vince Bielinski |
| 35 | L0 | DE | DE_Venus | Venus YFP | | Bernardo Pollak |
| 36 | L0 | DE | DE_3xStop | 3xStop codon - small part | | Bernardo Pollak |
| 37 | L0 | DE | DE_peroxTP | Pt Peroxisomal targetting peptide | | Chris Dupont |
| 38 | L0 | EF | EF_PtBle | ShBle with FcpF promoter | | Vince Bielinski |
| 39 | L0 | EF | EF_B0015 | BBa_B0015 | L0>L1 Rx control | iGEM |
| 40 | L0 | EF | EF_t49202 | Pt-t49202 | | Patrick Brunson |
| 41 | L0 | EF | EF_tFcpB | Pt-tFcpB | | Patrick Brunson |
| 42 | L0 | EF | EF_tH4 | Pt-tH4 | | Patrick Brunson |
| 43 | L0 | EF | EF_tNR | Pt-tNR | | Patrick Brunson |
| 44 | L0 | EF | EF_tUbq3 | Polyubiquitin 3 terminator | | plasmid [12] |
| 45 | L0 | EF | EF_tNos-t35S | nosT-35ST double terminator | | Bernardo Pollak [3] |
| 46 | L0 | EF | EF_t35S | CaMV 35S terminator | | Bernardo Pollak/[13] |
| 47 | L0 | EF | EF_thsp18.2(GB0035) | Hsp18.2 terminator | | GB 2.0 kit |
| 48 | L0 | EF | EF_tADH1 | tADH1 terminator | SpectR; Chemical synthesis | Matute & Núñez |
| 49 | L0 | EF | EF_tCYC1 | tCYC1 terminator | SpectR; Domestication from p426 (Addgene #43803; [7] ). | Matute & Núñez/[7] |
| 50 | L0 | * | Venus | Venus FP | | Bernardo Pollak/[14] |
| 51 | L0 | * | mTurquoise2 | mTurquoise2 cyan FP | | Bernardo Pollak/[15] |
| 52 | L0 | * | mTagBFP2 | mTagBFP2 blue FP | | Bernardo Pollak/[9] |
| 53 | L0 | * | Lti6b-Tag | Lti6b Membrane targetting peptide | | Bernardo Pollak/[16] |
| 54 | L0 | * | N7 | N7 Nuclear targetting peptide | | Bernardo Pollak/[16] |

**Supplementary Table S10. L0 part list.**

| N | Level | Pos. | Name | Description | P1 | P2 | P3 | P4 | P5 | Rec |
|---|---|---|---|---|---|---|---|---|---|---|
| 1 | L1 | 1 | pCAL1-1_PTCv2 | Conjugation and prop. Elements | AC_CEN_ARS_HIS | CE_OriTv2 | EF_PtBle | | | pCAo-1 |
| 2 | L1 | 2 | pCAL1-2_pNR-Venus-PeroxTP-tNR | pNR driven peroxisome Venus | AC_pNR | CD_Venus | DE_peroxTP | EF_tNR | | pCAo-2 |
| 3 | L1 | 3 | pCAL1-3_pH4-MitoTP-mTurquoise2-nH4 | pH4 driven mitochondrial mTurquoise2 | AC_pH4 | CD_MitoTP | DE_mTurquoise2 | EF_tH4 | | pCAo-3 |
| 4 | L1 | 4 | pCAL1-4_p49202-mTagBFP2-t49202 | p49202 driven cytoplasmic mTagBFP2 | AC_p49202 | CD_mTagBFP2 | DE_3xStop | EF_t49202 | | pCAo-4 |
| 5 | L1 | 1 | pCAL1-1_pUBQ10-Venus-Lti6b-Ubq3T | pUBQ10 driven membrane Venus | AC_pUBQ10 | Venus | Lti6b-Tag | EF_tUbq3 | | pCAo-1 |
| 6 | L1 | 2 | pCAL1-2_pUBQ10-mTurquoise2-N7-nosT-35ST | pUBQ10 driven nuclear mTurquoise2 | AC_pUBQ10 | mTurquoise2 | N7 | EF_tNos-t35S | | pCAo-2 |
| 7 | L1 | 3 | pCAL1-3_pUBQ10-mRuby3-35ST | pUBQ10 driven cytoplasmic mRuby3 | AC_pUBQ10 | CE_mRuby3 | EF_t35S | | | pCAo-3 |
| 8 | L1 | 4 | pCAL1-4_pUBQ10-mTagBFP2-Lti6B - THsp18.2 | pUBQ10 driven membrane mTagBFP2 | AC_pUBQ10 | mTagBFP2 | Lti6b-Tag | EF_thsp18.2(GB0035) | | pCAo-4 |
| 9 | L1 | 1 | pSBL1-1_pUBQ10-Venus-Lti6b-Ubq3T | pUBQ10 driven membrane Venus | AC_pUBQ10 | Venus | Lti6b-Tag | EF_tUbq3 | | pSBo-1 |
| 10 | L1 | 2 | pSBL1-2_pUBQ10-mTurquoise2-N7-nosT-35ST | pUBQ10 driven nuclear mTurquoise2 | AC_pUBQ10 | mTurquoise2 | N7 | EF_tNos-t35S | | pSBo-2 |
| 11 | L1 | 3 | pSBL1-3_pUBQ10-mRuby3-35ST | pUBQ10 driven cytoplasmic mRuby3 | AC_pUBQ10 | CE_mRuby3 | EF_t35S | | | pSBo-3 |
| 12 | L1 | 4 | pSBL1-4_pUBQ10-mTagBFP2-Lti6B - THsp18.2 | pUBQ10 driven membrane mTagBFP2 | AC_pUBQ10 | mTagBFP2 | Lti6b-Tag | EF_thsp18.2(GB0035) | | pSBo-4 |
| 13 | L1 | 1 | pCAL1-1_2u | L1-1 2u ori (yeast) | AF_2u | - | - | - | - | pCAo-1 |
| 14 | L1 | 1 | pCAL1-1_CEN | L1-1 CEN ori (yeast) | AF_CEN | - | - | - | - | pCAo-1 |
| 15 | L1 | 2 | pCAL1-2_pTDH3-mTagBFP2-tCYC1 | L1-2 pTDH3 encodes mTagBFP2 | AC_pTDH3 | CD_mTagBFP2 | DE_3xStop | EF_tCYC1 | - | pCAo-2 |
| 16 | L1 | 2 | pCAL1-2_pTDH3-mTurq2-tCYC1 | L1-2 pTDH3 encodes mTurquoise2 | AC_pTDH3 | CD_mTurquoise2 (yeast) | DE_3xStop | EF_tCYC2 | - | pCAo-2 |
| 17 | L1 | 2 | pCAL1-2_pTEF1-LuxR-tADH1 | L1-2 pTEF1 encodes LuxR | AC_pTEF1 | CD_LuxR | DE_3xStop | EF_tADH1 | - | pCAo-2 |
| 18 | L1 | 4 | pCAL1-4_Spacer2 | L1-4 Spacer2 | AF_Spacer2 | - | - | - | - | pCAo-4 |
| 19 | L1 | 3 | pCAL1-3_URA3 | L1-3 URA3 | AF_URA3 | - | - | - | - | pCAo-3 |
| 20 | L1 | 1 | pCAL1-1_pLac-54-BeRFP-B0015 | L1-1 pLac encodes mBeRFP | AB_R0010 | BC_RiboJ54 | CD_mBeRFP | DE_3xStop | EF_B0015 | pCAo-1 |
| 21 | L1 | 2 | pCAL1-2_pLac-54-YFP-B0015 | L1-2 pLac encodes YFP | AB_R0010 | BC_RiboJ54 | CD_ E0030 (YFP) | DE_3xStop | EF_B0015 | pCAo-2 |
| 22 | L1 | 3 | pCAL1-3_pLac-54-mTagBFP2-B0015 | L1-3 pLac encodes mTagBFP2 | AB_R0010 | BC_RiboJ54 | CD_mTagBFP2 | DE_3xStop | EF_B0015 | pCAo-3 |
| 23 | L1 | 4 | pCAL1-4_pLac-54-sfGFP-B0015 | L1-4 pLac encodes sfGFP | AB_R0010 | BC_RiboJ54 | CD_sfGFP | DE_3xStop | EF_B0015 | pCAo-4 |
| 24 | L1 | 4 | pCAL1-4_pLac-54-turq2-B0015 | L1-4 pLac encodes mTurquoise2 | AB_R0010 | BC_RiboJ54 | CD_Tur2 | DE_3xStop | EF_B0015 | pCAo-4 |
| 25 | L1 | 1 | pSBL1-1_pLac-54-BeRFP-B0015 | L1-1 pLac encodes mBeRFP | AB_R0010 | BC_RiboJ54 | CD_mBeRFP | DE_3xStop | EF_B0015 | pSBo-1 |
| 26 | L1 | 2 | pSBL1-2_pLac-54-YFP-B0015 | L1-2 pLac encodes YFP | AB_R0010 | BC_RiboJ54 | CD_ E0030 (YFP) | DE_3xStop | EF_B0015 | pSBo-2 |
| 27 | L1 | 3 | pSBL1-3_pLac-54-mTagBFP2-B0015 | L1-3 pLac encodes mTagBFP2 | AB_R0010 | BC_RiboJ54 | CD_mTagBFP2 | DE_3xStop | EF_B0015 | pSBo-3 |
| 28 | L1 | 4 | pSBL1-4_pLac-54-sfGFP-B0015 | L1-4 pLac encodes sfGFP | AB_R0010 | BC_RiboJ54 | CD_sfGFP | DE_3xStop | EF_B0015 | pSBo-4 |
| 30 | L1 | 4 | pSBL1-4_pLac-54-turq2-B0015 | L1-4 pLac encodes mTurquoise2 | AB_R0010 | BC_RiboJ54 | CD_Tur2 | DE_3xStop | EF_B0015 | pSBo-4 |
| 31 | L1 | 3 | pCAL1-3_Spacer 1 | L1-3 Spacer 1 | AF_Spacer1 | - | - | - | - | pCAo-3 |
| 32 | L1 | 2 | pCAL1-2_p49202-LuxR-t49202 | L1-2 p49202 encodes LuxR | AC_p49202 | CD_LuxR | DE_3xStop | EF_t49202 | - | pCAo-2 |
| 33 | L1 | 2 | pCAL1-2_pH4-mTagBFP2-tH4 | L1-2 pH4 encodes mTagBFP2 | AC_pH4 | CD_mTagBFP2 | DE_3xStop | EF_tH4 | - | pCAo-2 |
| 34 | L1 | 1 | pL1-1_35SmR3 | L1-1_35SmR3 | CaMV35S | mRuby3 | nosT | | - | pOdd-1 |
| 35 | L1 | 2 | pL1-2_35SmT2 | L1-2_35SmT2 | CaMV35S | mTurquoise2 | N7 | nosT | - | pOdd-2 |
| 36 | L1 | 3 | pL1-3_35SVe | L1-3_35SVe | CaMV35S | Venus | N7 | nosT | - | pOdd-3 |
| 37 | L1 | 4 | pL1-4_35SmR3 | L1-4_35SmR3 | CaMV35S | mRuby3 | nosT | | - | pOdd-4 |
| 38 | L1 | 4 | pCAL1-4_sfGFP | L1 assembly sfGFP test cassette | AB_J23101 | BC-B0034m | CE_sfGFP | EF_B0015 | - | pCAo-4 |
| 39 | L1 | 4 | pCOL1-4_sfGFP | L1 assembly sfGFP test cassette | AB_J23101 | BC-B0034m | CE_sfGFP | EF_B0015 | - | pCOo-4 |
| 40 | L1 | 4 | pSBL1-4_sfGFP | L1 assembly sfGFP test cassette | AB_J23101 | BC-B0034m | CE_sfGFP | EF_B0015 | - | pSBo-4 |
| 41 | L1 | 4 | pANL1-4_sfGFP | L1 assembly sfGFP test cassette | AB_J23101 | BC-B0034m | CE_sfGFP | EF_B0015 | - | pANo-4 |

**Supplementary Table S11. L1 assembly list.**

| N | Level | Pos. | Name | Description | P1 | P2 | P3 | P4 | Rec |
|---|---|---|---|---|---|---|---|---|---|
| 1 | L2 | 1 | pCAL2-1_4xFP | L2-1 Fluorescent protein reporters 4x (plant) | pCAL1-1_pUBQ10-Venus-LTi6b | pCAL1-2_pUBQ10-mTurquoise2-N7 | pCAL1-3_pUBQ10-mRuby3 | pCAL1-4_pUBQ10-mTagBFP2-LTi6B | pCAe-1 |
| 2 | L2 | 1 | pSBL2-1_4xFP | L2-1 Fluorescent protein reporters 4x (plant) | pSBL1-1_pUBQ10-Venus-LTi6b | pSBL1-2_pUBQ10-mTurquoise2-N7 | pSBL1-3_pUBQ10-mRuby3 | pSBL1-4_pUBQ10-mTagBFP2-LTi6B | pSBe-1 |
| 3 | L2 | 1 | pCAL2-1_Yeast-NF | L2-1 pTEF1 encodes LuxR (yeast) | pCAL1-1_CEN | pCAL1-2_pTEF1-LuxR-tADH1 | pCAL1-3_URA3 | pCAL1-4_Spacer2 | pCAe-1 |
| 4 | L2 | 1 | pCAL2-1_Yeast-mT2 | L2-1 Fluorescent reporter mTurquoise2 (yeast) | pCAL1-1_2u | pCAL1-2_pTDH3-mTurq2-tCYC1 | pCAL1-3_URA3 | pCAL1-4_Spacer2 | pCAe-1 |
| 5 | L2 | 1 | pSBL2-1_Yeast-B | L2-1 Fluorescent reporter mTagBFP2 (yeast) | pCAL1-1_2u | pCAL1-2_pTDH3-mTagBFP2-tCYC1 | pCAL1-3_URA3 | pCAL1-4_Spacer2 | pSBe-1 |
| 6 | L2 | 1 | pCAL2-1_RYBG | L2-1 Fluorecent protein reporters (E.coli) | pCAL1-1_pLac-54-BeRFP-B0015 | pCAL1-2_pLac-54-YFP-B0015 | pCAL1-3_pLac-54-mTagBFP2-B0015 | pCAL1-4_pLac-54-sfGFP-B0015 | pCAe-1 |
| 7 | L2 | 1 | pCAL2-1_RYBmT2 | L2-1 Fluorecent protein reporters (E.coli) | pCAL1-1_pLac-54-BeRFP-B0015 | pCAL1-2_pLac-54-YFP-B0015 | pCAL1-3_pLac-54-mTagBFP2-B0015 | pCAL1-4_pLac-54-turq2-B0015 | pCAe-1 |
| 9 | L2 | 1 | pSBL2-1_RYBG | L2-1 Fluorecent protein reporters (E.coli) | pSBL1-1_pLac-54-BeRFP-B0015 | pSBL1-2_pLac-54-YFP-B0015 | pSBL1-3_pLac-54-mTagBFP2-B0015 | pSBL1-4_pLac-54-sfGFP-B0015 | pSBe-1 |
| 10 | L2 | 1 | pSBL2-1_RYBmT2 | L2-1 Fluorecent protein reporters (E.coli) | pSBL1-1_pLac-54-BeRFP-B0015 | pSBL1-2_pLac-54-YFP-B0015 | pSBL1-3_pLac-54-mTagBFP2-B0015 | pSBL1-4_pLac-54-turq2-B0015 | pSBe-1 |
| 12 | L2 | 1 | pSBL2-1_Pt-B | L2-1 Pt Fluorescent reporter mTagBFP2 | pCAL1-1_PTCv2 | pCAL1-2_pH4-mTagBFP2-tH4 | pCAL1-3_Spacer 1 | pCAL1-4_Spacer2 | pSBe-1 |
| 13 | L2 | 1 | pCAL2-1_Pt-NF | L2-1 Pt p49202 encodes LuxR | pCAL1-1_PTCv2 | pCAL1-2_p49202-LuxR-t49202 | pCAL1-3_Spacer 1 | pCAL1-4_Spacer2 | pCAe-1 |
| 14 | L2 | 1 | pCAL2-1_FPrep | L2-1 Pt multi-spectral reporter | pCAL1-1_PTCv2 | pCAL1-2_pNR-Venus-PeroxTP | pCAL1-3_pH4-MitoTP-mTurquoise2 | pCAL1-4_p49202-mTagBFP2 | pCAe-1 |
| 15 | L2 | 1 | pL2-1_all | L2-1 encodes 4 plant FP reporters | pL1-1_35SmR3 | pL1-2_35SmT2 | pL1-3_35SVe | pL1-4_35SmR3 | pEven-1 |
| 16 | L2 | 2 | pL2-2_all | L2-2 encodes 4 plant FP reporters | pL1-1_35SmR3 | pL1-2_35SmT2 | pL1-3_35SVe | pL1-4_35SmR3 | pEven-2 |
| 17 | L2 | 3 | pL2-3_all | L2-3 encodes 4 plant FP reporters | pL1-1_35SmR3 | pL1-2_35SmT2 | pL1-3_35SVe | pL1-4_35SmR3 | pEven-3 |
| 18 | L2 | 4 | pL2-4_all | L2-4 encodes 4 plant FP reporters | pL1-1_35SmR3 | pL1-2_35SmT2 | pL1-3_35SVe | pL1-4_35SmR3 | pEven-4 |
| 19 | L2 | 4 | pCAL2-4_sfGFP | 4 TU construct with sfGFP cassette | pL1-1_35SmR3 | pL1-2_35SmT2 | pL1-3_35SVe | pCAL1-4_sfGFP | pCAe-4 |
| 20 | L2 | 4 | pCOL2-4_sfGFP | 4 TU construct with sfGFP cassette | pL1-1_35SmR3 | pL1-2_35SmT2 | pL1-3_35SVe | pCAL1-4_sfGFP | pCOe-4 |
| 21 | L2 | 4 | pSBL2-4_sfGFP | 4 TU construct with sfGFP cassette | pL1-1_35SmR3 | pL1-2_35SmT2 | pL1-3_35SVe | pCAL1-4_sfGFP | pSBe-4 |
| 22 | L2 | 4 | pANL2-4_sfGFP | L2 assembly test construct with sfGFP cassette | pL1-1_35SmR3 | pL1-2_35SmT2 | pL1-3_35SVe | pCAL1-4_sfGFP | pANe-4 |
| 23 | L3 | 4 | pCAL3-4_sfGFP | 16 TU construct with sfGFP cassette | pL2-1_all | pL2-2_all | pL2-3_all | pCAL2-4_sfGFP | pCAo-4 |
| 24 | L3 | 4 | pCOL3-4_sfGFP | 16 TU construct with sfGFP cassette | pL2-1_all | pL2-2_all | pL2-3_all | pCAL2-4_sfGFP | pCOo-4 |
| 25 | L3 | 4 | pSBL3-4_sfGFP | 16 TU construct with sfGFP cassette | pL2-1_all | pL2-2_all | pL2-3_all | pCAL2-4_sfGFP | pSBo-4 |
| 26 | L3 | 4 | pANL3-4_sfGFP | 16 TU construct with sfGFP cassette | pL2-1_all | pL2-2_all | pL2-3_all | pCAL2-4_sfGFP | pANo-4 |
| 27 | L3 | 1 | pCAL3-1_allx4 | 16 TU test assembly construct | pL2-1_all | pL2-2_all | pL2-3_all | pL2-4_all | pCAo-1 |
| 28 | L3 | 2 | pCAL3-2_allx4 | 16 TU test assembly construct | pL2-1_all | pL2-2_all | pL2-3_all | pL2-4_all | pCAo-2 |
| 29 | L3 | 3 | pCAL3-3_allx4 | 16 TU test assembly construct | pL2-1_all | pL2-2_all | pL2-3_all | pL2-4_all | pCAo-3 |
| 30 | L4 | - | L4-4_sfGFP | 64 TU 4 piece assembly | pCAL3-1_allx4 | pCAL3-2_allx4 | pCAL3-3_allx4 | pCAL3-4_sfGFP | - |

**Supplementary Table S12. L2, L3 and L4 assembly list**

#### Plate fluorometry data analysis

Dynamical measurements of bacterial growth (OD600 absorbance) and fluorescence obtained by plate reader fluorometry were analysed with python routines included in Jupyter notebooks described below.
**load-data.ipynb :** was used to load and format the data from the plate reader to the experimental database.
**data-treatment.ipynb:** was used to subtract absorbance and fluorescence background values.
**data-analysis-uLoop.ipynb:** was used to organise data, apply models to obtain relevant parameters, perform the statistics and make the plots.
Please read the comments in the code for further details [17].
Signal readings ($I_m$) were cleaned by subtracting the corresponding background ($I_b$) values in each case as shown in equation 1.

$$I_i(t) = I_m(t) + I_b(t). \tag{1}$$

Next, cross measurements of red and green fluorescence from GFP and RFP control cultures, respectively, were performed to determine any bleed-through signal. Red and green fluorescence signals were measured from cell cultures containing pCAL1-4-sfGFP plasmid and from AKR1 cell cultures, respectively. Empirical functions were fitted to each data set as shown in Supplementary Figures S19 and Figures S20, which relate bleed-through fluorescent signal with growth (OD600). These functions were applied to each data series to subtract the proper background signals (Figures S21 and Figures S22). Relevant parameters related to fluorescent expression were extracted from pre-treated data. As sfGFP and RFP genes were under constitutive expression, they were expected to display a constant fluorescent ratio. In accordance to this, sfGFP signal followed roughly a linear relation with respect to mRFP1 signal (Supplementary Figures S6-S10). Next, a linear fit was performed to characterise each data set as the expression ratio obtained from the slope of these linear functions (Supplementary Figures S6-S10). Mean ratio values were computed by vector set and assembly level over the 3 whole replicates (N=3 and 6 technical replicates of each day)(Figure 4A and Supplementary table S1). To measure the variability of expression for each vector set, the coefficient of variation (standard deviation divided by the mean reported as a percentage) was computed for each group (Supplementary table S7).

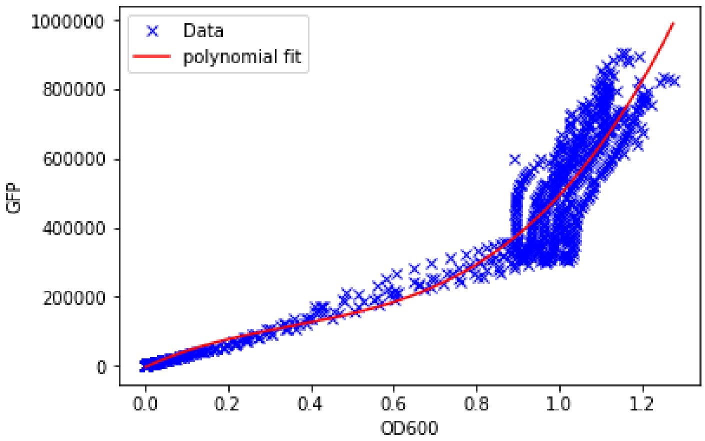

**Supplementary Figure S19. AKR1 green bleed-trough estimation.**
An empirical function was fitted to GFP (ex: 485 em: 516) vs OD600 from AKR1 control cultures (N = 3). Blue crosses correspond to each data point and the red line represents the fitted polynomial function.

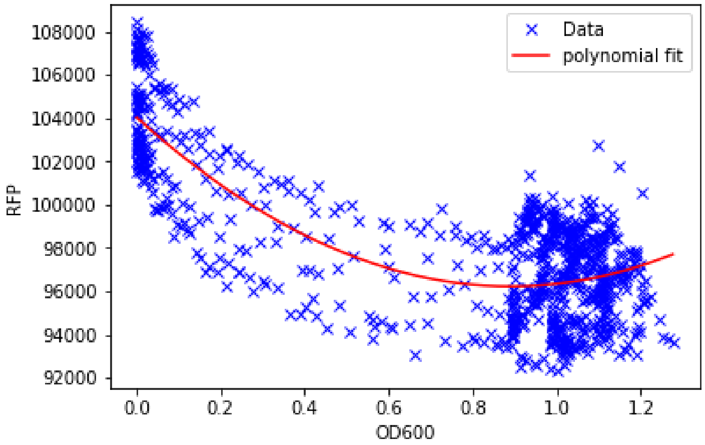

**Supplementary Figure S20. Red fluorescence bleed-trough estimation.**
An empirical function was fitted to RFP (ex: 585 em: 620) vs OD600 from AKR1 control cultures (N = 3). Blue crosses correspond to each data point and the red line represents the fitted polynomial function.

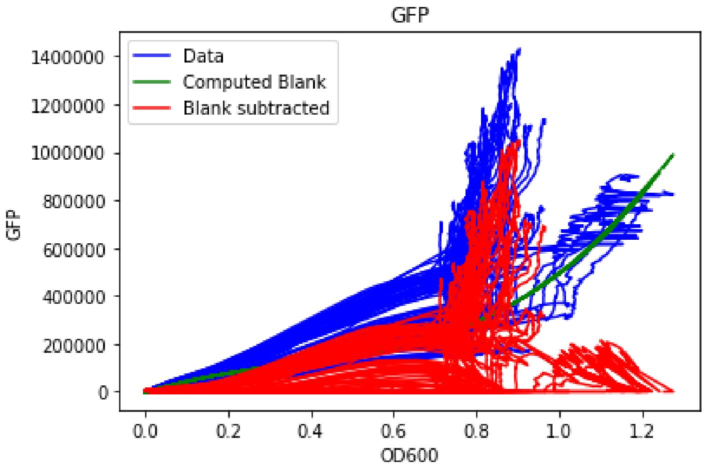

**Supplementary Figure S21. GFP background subtraction.**
A fitted function was used to remove the background from GFP measurements for each data set. Blue lines correspond to raw GFP values, the green line corresponds to the empirical background function and red lines represent background-subtracted data.

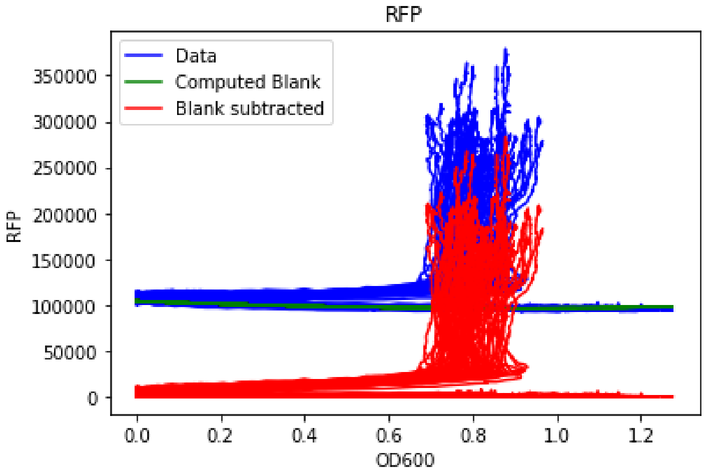

**Supplementary Figure S22. RFP background subtraction.**
A fitted function was used to remove the background from RFP measurements for each data set. Blue lines correspond to raw RFP values, the green line represents the empirical background function and red lines correspond to background-subtracted data.

#### Growth rate measurements

First, blank wells containing M9-Glycerol medium only were measured to obtain absorbance background values. As the OD600 blank values were small and approximately constant, a single mean was taken and used as a background value (OD600 blank value = 0.081 units) and later subtracted to all the other well values. Next, the effect of different plasmid kits and levels on cell growth was analysed by fitting a Gompertz model [18] over OD600 absorbance data for each well, according to equation 2:

$$ln\left[\frac{A}{A_0}\right] = k \cdot exp\left\{ - exp\left[\frac{\mu_{max} \cdot e}{k} \cdot (\lambda - t) + 1\right]\right\} \tag{2}$$

The maximum growth rate parameter ($\mu_{max}$) was obtained for each data series. They were grouped and analysed by level and vector kit as shown in Figure 4C and supplementary Table S3.
